# Glutamate receptor composition at *Drosophila* neuromuscular junctions depends on developmental stage and muscle identity

**DOI:** 10.64898/2026.03.17.712218

**Authors:** Anne Sustar, Chengjie Qiu, Yu Xiong, Dion Dickman, John C. Tuthill

## Abstract

The neuromuscular junction (NMJ) of larval *Drosophila* is widely used for studying synaptic transmission. Larval body wall muscles express five ionotropic glutamate receptor (iGluR) subunits that assemble into two tetrameric complexes, with subunit composition determining the strength and plasticity of synaptic transmission. Because NMJ function has been extensively characterized in larvae, it has commonly been assumed that adult fly NMJs share this molecular composition, despite substantial differences between life stages. Here, we systematically compare glutamate receptor expression across larval and adult *Drosophila* muscles. We find that adult leg and flight muscles exhibit different iGluR expression than larvae, lacking expression of several receptors previously considered essential. Adjacent muscles within the adult femur express distinct iGluRs, suggesting specialization of flexor and extensor muscles. Finally, the glutamate-gated chloride channel (GluClα) is expressed extrasynaptically in adult but not larval muscle fibers. Our results reveal unexpected heterogeneity in glutamate receptor expression across muscles and developmental stages, demonstrating the need for muscle-specific analyses in flies and other animals.

## Introduction

Body movement relies on the release of chemical neurotransmitters from motor neurons onto muscles at the neuromuscular junction (NMJ). In vertebrate animals, most motor neurons release the excitatory neurotransmitter acetylcholine (Dale et al., 1936), which acts on nicotinic receptors to elicit contraction of skeletal muscle. In contrast, most invertebrate motor neurons release glutamate (Takeuchi and Takeuchi, 1964, 1963), which binds to ionotropic glutamate receptors (iGluRs) expressed in muscle (Schuster et al., 1991). However, muscles are not uniform effectors; they are highly specialized to control distinct body parts and behaviors ranging from walking to flight (Hooper and Thuma, 2005; Schiaffino et al., 2025). Given this functional specialization, how is the expression and subunit composition of postsynaptic neurotransmitter receptors regulated to support the function of diverse muscles? Determining how receptors are tailored to the specific biophysical demands of individual muscles, and how this tuning shifts during development, is essential for understanding the molecular mechanisms that underlie adaptive motor control.

A powerful experimental system to investigate molecular, developmental, and physiological mechanisms of neuromuscular signaling is the larval NMJ of the fruit fly, *Drosophila*, which enables high-resolution analysis of identified motor neurons and muscles with electrophysiology and immunohistochemistry (He and Dickman, 2025). Flies, like humans, encode 16 GluR genes **(Figure 1A**; Li et al., 2016). Five ionotropic receptor (iGluR) subunits are expressed in larval muscle, where two subtypes mediate excitatory synaptic transmission (DiAntonio, 2006; Thomas and Sigrist, 2012). Similar to vertebrate

**Figure 1.**
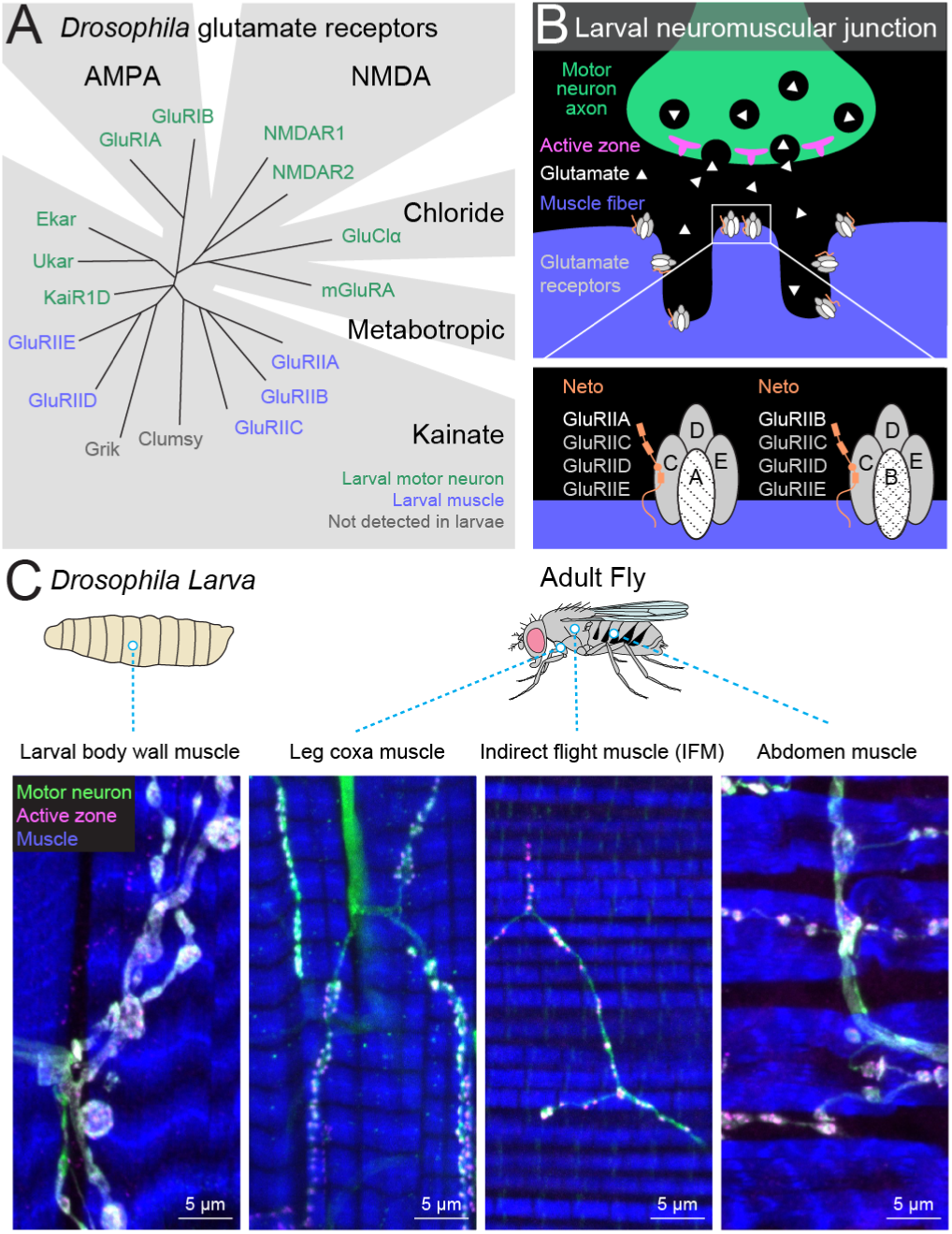
Glutamate receptors at *Drosophila* neuromuscular junctions (NMJs). (A) Phylogeny of 16 glutamate receptors in the *Drosophila* genome and their reported expression at the larval NMJ (based on Li et al., 2016). (B) Schematic of the larval *Drosophila* fly NMJ (top) and composition of glutamate receptors (bottom). Larval GluRs occur as two distinct tetrameric complexes, each with the same auxiliary subunit (Neto). (C) Example fluorescence images of larval and adult NMJs. Motor neurons are labeled by driving GFP with vGlut-Gal4 (green). Active zones are labeled with an antibody against Brp (Wagh et al., 2026) (magenta). Muscle is labeled with phalloidin (blue).

AMPA-type receptors, fly iGluRs function as tetrameric complexes that require three obligate subunits (GluRIIC, GluRIID, and GluRIIE) along with a fourth, either GluRIIA or GluRIIB (**Figure 1B**; Han et al., 2024, 2015). The specific subunit composition dictates synaptic efficacy: GluRIIA-containing receptors generate large, stable synaptic currents, while GluRIIB complexes desensitize rapidly (DiAntonio, 2006; Thomas and Sigrist, 2012). Synaptic transmission at the larval NMJ is also regulated homeostatically: when postsynaptic GluRs are genetically or pharmacologically suppressed, the presynaptic motor neuron increases neurotransmitter release to maintain stable postsynaptic responses (He and Dickman, 2025). Distinct from this excitatory machinery, the inhibitory glutamate-gated chloride channel (GluClα) functions presynaptically in larval motor neurons to mediate presynaptic homeostatic depression, a mechanism that adaptively dampens neurotransmitter release at the NMJ (Li et al., 2021). Due to the extensive characterization of these mechanisms in the larva, we and others assumed that this canonical molecular architecture is preserved in the adult fly. Only one pioneering study from Rivlin et al. (2004) observed that two larval receptor subunits (GluRIIA/GluRIIB) cluster at only a subset of thoracic muscles, hinting at differences in receptor expression between larvae and adults.

Compared to the NMJ of *Drosophila* larva, the physiology and molecular architecture of adult fly NMJs remain far less explored. Physiological recordings have been made from adult flight and proboscis muscles (Mahoney et al., 2014; Trimarchi and Murphey, 1997), but these studies address other questions and did not characterize the molecular composition of the NMJ. This gap is significant because the biomechanical context of movement changes fundamentally during metamorphosis (Agrawal and Tuthill, 2022). Larval body wall muscles mediate slow, rhythmic peristaltic locomotion by pulling against a hydraulic skeleton. In contrast, adult muscles drive a rigid exoskeleton to produce rapid behaviors such as flight and walking. Given these varied biophysical demands, it may be advantageous for specialized muscles to adapt the molecular composition of their synapses. Furthermore, with the emergence of comprehensive connectomes of the adult fly nervous system (Bates et al., 2025; Berg et al., 2025) and biomechanical body models (Vaxenburg et al., 2025; Wang-Chen et al., 2024), defining the molecular identity of neuromuscular synapses is a critical missing link required for integrative modeling of how the nervous system actuates the body.

Here, using genetic reporter lines, immunohistochemistry (IHC), and analysis of single-cell RNA sequencing data, we find that GluR expression in *Drosophila* is substantially more diverse than previously appreciated. We find that the receptor landscape in adult leg and flight muscles diverges substantially from the larval pattern; unexpectedly, we did not detect several of the GluR subunits that are required for glutamate receptor function at the larval NMJ. In contrast, adult abdominal muscles retain the canonical larval receptor complement, consistent with the fact that the larval musculature is transformed into the abdominal musculature during metamorphosis (Currie and Bate, 1991; Kimura and Truman, 1990). We also identify an extrasynaptic population of glutamate-gated chloride channels (GluClα) specific to adult muscle, which may account for the hyperpolarizing glutamate responses described in classical studies of other insects (Cull-Candy, 1976; Dudel et al., 1989; Lea and Usherwood, 1973). Collectively, these results reveal a diversity in NMJ composition, suggesting that synaptic composition is specialized to match the functional requirements of specific muscles.

## Results

### Larval and adult muscles express distinct GluRs

Consistent with past literature, GAL4 expression patterns and antibody labeling confirmed that all five GluRII subunits are expressed at NMJs throughout larval body wall muscle (**Figures 2C, s1**). We observed a similar expression pattern in the adult abdomen (**Figure 2C, s2**), although the labeling of *GluRIIB* and *GluRIIE* was weaker than in the larva. As expected, all larval and adult muscles expressed *Neto-β*, the primary auxiliary subunit required for iGluR receptor trafficking and function (**Figure 2D**). Expression of other Kainate-type receptors (*Grik*, *KaiR1D*, *Ukar*, *Ekar*), as well as AMPA, NMDA, and the sole metabotropic GluR were not detected in larval or adult muscles.

**Figure 2.**
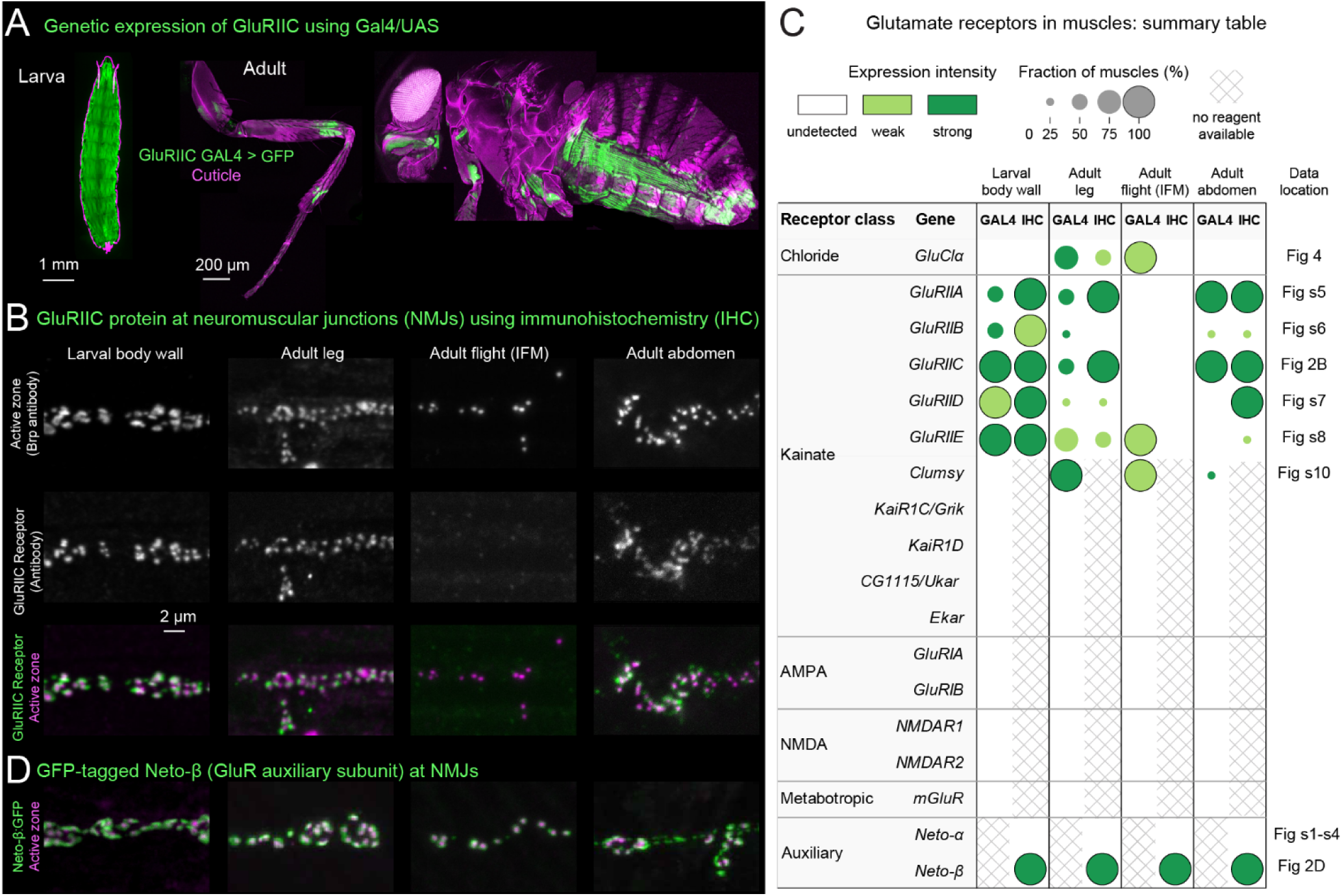
GluR subunit expression mapping in larval and adult muscles. (A) Representative confocal images showing GFP expression driven by a GAL4 reporter line (GluRIIC-GAL4) in larval and adult muscles. (B) Representative images showing antibody staining of GluRIIC at larval and adult neuromuscular junctions. (C) Summary table of Gal4 expression/antibody staining for all GluRs in larval and adult muscles. Note that antibodies and tagged proteins are combined in a single immunohistochemistry (IHC) column. Note that this initial screen focused on the coxa segment of the adult leg. See supplemental figures for representative images and methods for a complete list of reagents screened. (D) Representative images showing the GluR auxiliary subunit *Neto-β* tagged with GFP at larval and adult neuromuscular junctions.

Surprisingly, in adult indirect flight muscles we did not detect any of the five GluRII subunits by antibody staining or endogenous tagging, and GAL4 reporters for these subunits were either undetectable or, in the case of GluRIIE, extremely faint (**Figures 2C, s2-s3, s5-s8;** see Discussion for our interpretation of this discrepancy). Leg coxa muscles expressed subsets of GluRII subunits that varied across specific muscle subtypes (**Figure 2C, s5-8**), though this expression was consistent across flies (**Figure s9**). In all tissues where IIA-E are expressed at the NMJ, receptors were localized directly adjacent to the active zone (Figure 2B, s5-8). Interestingly, the one iGluR that was broadly expressed throughout the adult (e.g., leg, proboscis, neck, haltere and indirect/direct flight muscles) was the kainate-type iGluR *Clumsy* (**Figures 2C, s10**). We did not detect *Clumsy* expression in larval muscles, and it was expressed only weakly in abdominal muscles. The *Clumsy* gene is phylogenetically related to GluRIIA/B/C (Li et al., 2016), but its function is poorly understood compared to the other GluRII subunits expressed in larval muscle.

### Different leg muscles express distinct iGluRs

Our screen revealed that the expression patterns of iGluR-specific Gal4 lines in the adult leg were often patchy: different muscles appeared to express different iGluR subunits (**Figures 2A, s5-s8**). These expression patterns were consistent across flies (**Figure s9**). To examine muscle-specific iGluR expression in more detail, we focused on the femur of the fly’s front leg (**Figure 3A-B**), for which we have previously characterized motor neuron physiology, muscle innervation, and muscle force production (Azevedo et al., 2020). Based on motor unit organization and tendon attachment, muscle fibers in the tibia can be grouped into four groups: a long tendon muscle, a tibia extensor, and two tibia flexors (Azevedo et al., 2024; Lesser et al., 2024).

**Figure 3.**
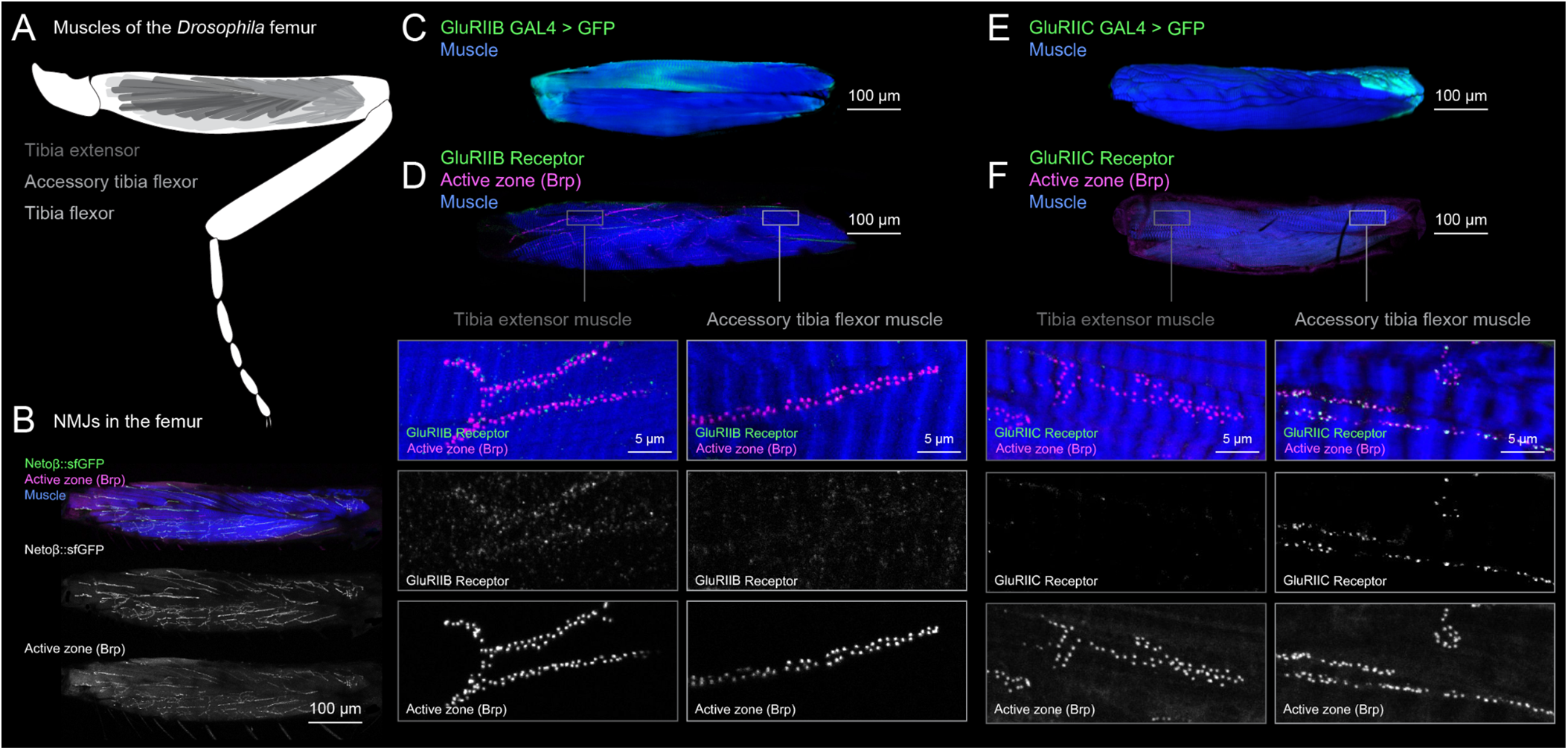
Differential GluRIIB vs GluRIIC subunit expression in adult muscle subtypes. (A) Schematic of the fly’s front leg with annotations of three key muscle groups in the femur. (B) Femur with Netoβ::sfGFP (green) in NMJs (Brp, magenta) in all femur muscle types (phalloidin, blue). (C) 3D rendering of a confocal stack of femur muscles (phalloidin, blue) and GluRIIB GAL4> UAS-GFP (green). GFP is expressed specifically in tibia extensor muscle fibers. (D) Representative antibody staining images showing presence of GluRIIB in a tibia extensor muscle (left) and no detectable GluRIIB in an accessory tibia flexor muscle (right). (E) 3D rendering of a confocal stack of femur muscles (phalloidin, blue) and GluRIIC GAL4> UAS-GFP (green). GFP is expressed specifically in accessory tibia flexor muscle fibers. (F) Representative antibody staining images showing no detectable GluRIIC in a tibia extensor muscle (left) and presence of GluRIIC in an accessory tibia flexor muscle (right).

To examine differences in iGluR expression between muscles, we compared Gal4 reporter expression and antibody labeling at NMJs in the femur for two GluR subunits: GluRIIB and GluRIIC. We chose these two subunits because their GAL4 expression was restricted to distinct muscle fibers (**Figure 3C, E**). We found that protein labeling was consistent with GAL4 expression: GluRIIB was expressed at NMJs of tibia extensor muscles but was not detected in other muscles, including the accessory tibia flexors (**Figure 3C-D**). In contrast, GluRIIC was expressed at NMJs of the more distal accessory tibia flexor muscles but was not detected in the more proximal muscles (**Figure 3E-F**). The distal accessory tibia flexor fibers are innervated by the slow motor neurons that control slow, postural movements of the tibia (Azevedo et al., 2020). Because these two antibodies label complementary sets of NMJs within the same bisected femur, processed and imaged together, the failure to detect a subunit in one muscle cannot be attributed to a general failure of that antibody in adult tissue.

We also observed differences in iGluR expression within a single muscle. For example, GluRIIB is expressed in proximal fibers of the tibia extensor but was not detected in the more distal fibers (**Figure 3C-D**). Prior work has shown that the more proximal fibers are innervated by the fast extensor motor neuron (FETi), while the more distal fibers are innervated by the slow extensor motor neuron (SETi; Azevedo et al., 2024). These differences in GluRIIB expression within the tibia extensor muscle may reflect specialization of fast vs. slow twitch muscle fibers.

Overall, our results show that different muscles in the fly leg express different ensembles of GluR subunits. We also confirmed that all femur NMJs express the obligatory GluR accessory protein Neto (**Figure 3B**), affirming that all femur motor neurons are glutamatergic, despite their otherwise unexpected receptor expression. Given what is known about larval GluR subunit composition, this implies that at least some adult GluRs are composed of non-canonical GluR subunits. Further, the difference in GluR expression is interesting because our prior work has demonstrated major differences in the physiology and force production of different muscle groups, and even different fibers within the same muscle (Azevedo et al., 2020). Thus, GluR subunit composition may be specialized to support diverse functions, even within a muscle.

### RNA-seq data are consistent with adult muscle expression patterns

We next analyzed transcriptomic data of muscle cells from an existing single-nuclei RNA-seq dataset of adult *Drosophila* tissues (Li et al., 2022). Consistent with our results from GAL4 reporters and antibody staining, transcription of all GluRII genes was relatively low in clusters annotated as adult leg (**Figures s11-s13**) and indirect flight muscle (**Figure s12, s13**), and only slightly higher in a cluster annotated as abdominal muscle. In comparison, and consistent with our results using GAL4 reporter lines, GluClα and Clumsy were both highly expressed in all three muscle clusters. One discrepancy was that *Ukar* and *Grik* had elevated transcription in multiple muscle clusters, which we did not observe with GAL4 lines. The Ukar GAL4 line had no detectable expression in any tissue, so this reporter line may not be functional. Overall, however, expression patterns from adult RNA-seq data were largely consistent with our main conclusions from GAL4 reporters and antibody staining.

### Motor neurons express a distinct set of GluR genes compared to muscle

To determine whether GluR localization at the neuromuscular junction could be due to presynaptic expression, we also screened for GluR expression in motor neurons. We expressed GFP under the control of GluR-specific GAL4 lines and examined whole-mount expression in the central nervous system, as well as axons projecting to the legs, indirect flight muscles, and abdomen (**Figure s14**). As expected, we found that motor neurons expressed GluClα, which mainly functions as an inhibitory neurotransmitter in the *Drosophila* central nervous system (Liu and Wilson, 2013). We also observed that motor neurons expressed a subset of kainate, AMPA, NMDA, and mGluR receptors that was nonoverlapping with the expression patterns we observed in muscle. Overall, these results confirm that the patterns of GluR expression we document at neuromuscular junctions are not due to expression in presynaptic motor neurons.

### Extrasynaptic glutamate-gated chloride channels in adult muscle

One surprising result from our survey of GluR expression was the broad expression of the glutamate-gated chloride channel, GluClα, in adult leg and flight muscles, but not in larval body wall or adult abdomen (**Figure 2C**). GluClα was also the GluR gene with the highest level of expression in leg muscles based on the single nuclei RNA-seq data (**Figure 4A**). Although GluClα has been shown to function presynaptically in larval motor neurons (Li et al., 2021), its presence and function in *Drosophila* muscle has not previously been investigated.

**Figure 4.**
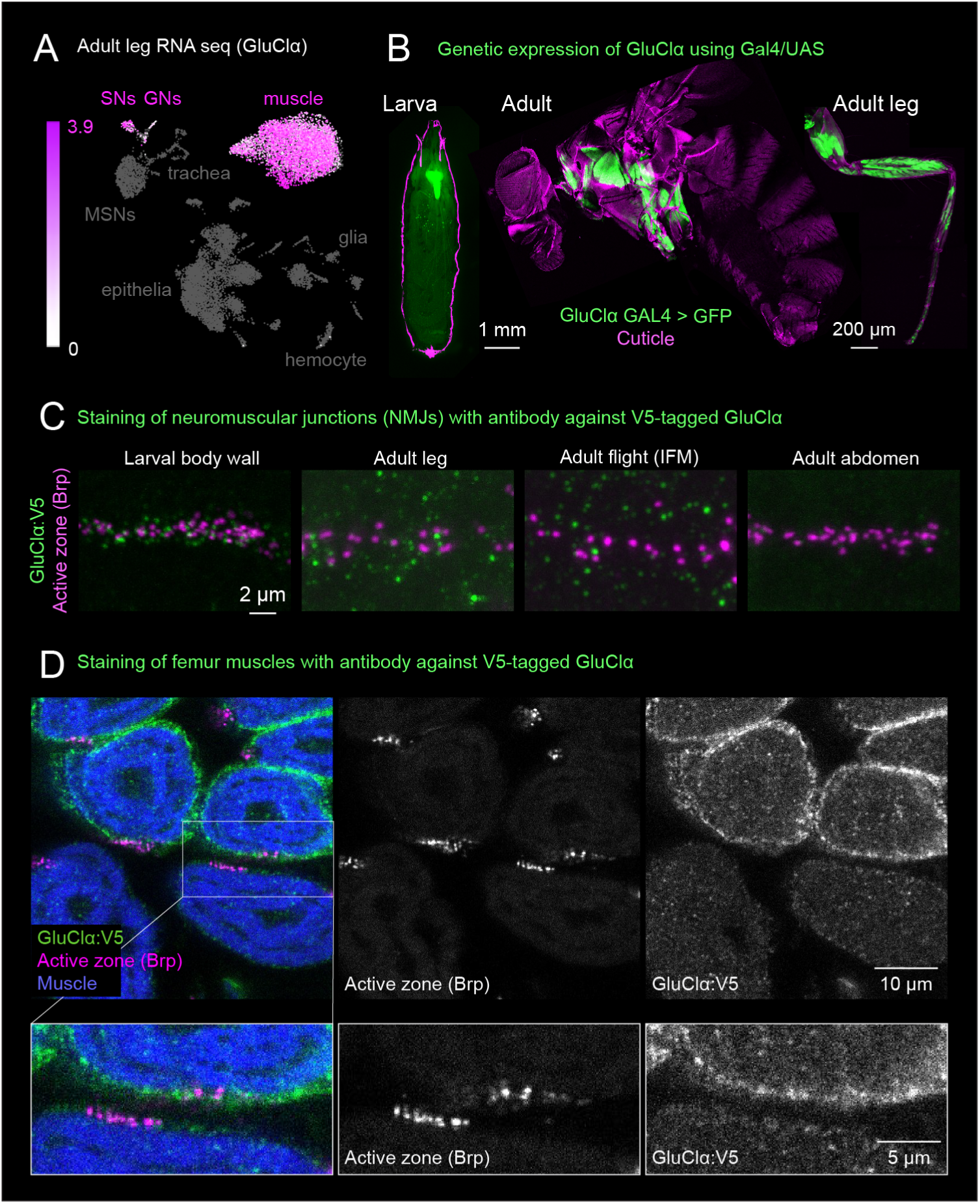
GluClα is expressed extrasynaptically in adult fly muscles. (A) Single-nuclei RNA-seq data shows strong expression of GluClα in adult leg muscles. (B) A Gal4 reporter for GluClα labels muscles in adult legs but not larval muscles. Expression in the larva is mainly in the nervous system, as expected. (C) Representative images showing antibody staining of endogenously-tagged GluClα:V5 in larval and adult muscles. Co-staining with an antibody against Brp reveals that GluClα does not co-localize with active zones. Labeling in larval muscle is likely from motor neurons (Li et al., 2021). (D) Labeling of GluClα protein with a V5 tag shows extrasynaptic expression throughout the muscle fiber, whereas active zones (magenta) are localized.

Whole-mount expression of GFP driven by two GluClα-Gal4 lines showed broad labeling of muscles throughout the flight and leg musculature (**Figure 4B**). However, using higher resolution imaging of an antibody against an endogenously-tagged GluClα:V5 protein (Sanfilippo et al., 2024), we found that puncta in leg and flight muscle did not cluster at synaptic active zones (**Figure 4C**). Rather, co-staining with phalloidin (**Figures 4D, s15**) revealed that the protein is distributed around the edges of the muscle fibers. This prominent V5 signal was not detected in wild-type negative controls **(Figure s15)**. Thus, unlike the other GluRs, GluClα localizes extrasynaptically in leg and indirect flight muscles. Although the function of extrasynaptic GluClα in muscle remains unclear, this result is consistent with classical electrophysiological studies in other invertebrates (e.g., locust, beetle, lobster) that described hyperpolarizing responses to extrasynaptic glutamate application (see Discussion).

## Discussion

In 2004, Rivlin and colleagues conducted a careful survey of NMJs in seven muscles in the adult *Drosophila* thorax. They established that adult NMJs are morphologically distinct from larval NMJs, and that GluRIIA and GluRIIB are expressed in muscle-specific subsets, with some muscles mysteriously lacking both. Importantly, they speculated that some adult muscles must be using different glutamate receptor subunits (which were unknown at the time) to build synapses. This problem has remained unexplored for the last two decades.

Here, we build on this work by surveying the expression of all 16 *Drosophila* glutamate receptor genes across larval and adult muscles. We find that glutamate receptor expression at *Drosophila* NMJs is considerably more heterogeneous than previously appreciated, with adult leg and flight muscles exhibiting receptor subunit combinations that diverge from the canonical patterns described in larvae. In adult indirect flight muscles we did not detect any of the five GluRII subunits required for larval NMJ function, suggesting that these specialized muscles may rely on alternative receptor combinations for neuromuscular transmission. Conversely, adult abdominal muscles mostly retain larval-like receptor expression, reflecting their shared developmental lineage. The identification of extrasynaptic GluClα in adult muscles establishes a possible mechanism for inhibitory glutamate responses described decades ago in other insects and suggests novel mechanisms for regulating muscle excitability.

### Adult leg and flight muscles lack expression of canonical larval glutamate receptors

Perhaps the most striking finding of our study is that some adult leg and flight muscles lack expression of the GluR subunits that have been characterized as essential for larval NMJ function. For example, we did not detect GluRIIA-E subunits in indirect flight muscles by any of three independent approaches: GAL4 reporter expression, antibody staining, and single-nucleus RNA-seq. These iGluR subunits were previously identified through genetic studies showing that mutations disrupting GluRIIC (Marrus et al., 2004), GluRIID (Featherstone et al., 2005), or GluRIIE (Qin et al., 2005) cause lethality in larvae, while double mutants lacking both GluRIIA and GluRIIB are similarly nonviable (DiAntonio et al., 1999; Petersen et al., 1997). Importantly, because these mutants die before metamorphosis, these experiments established a requirement for viability and for larval NMJ function, but could not test whether the same subunits are required at adult NMJs, which are formed during pupal development. Our observation that adult indirect flight muscles appear to function without detectable levels of these subunits suggests that the minimal receptor complement for glutamatergic neuromuscular transmission differs among NMJs within the same animal.

We found that Neto, the obligate auxiliary subunit required for GluR receptor function (Han et al., 2024; Kim et al., 2012), is expressed in all larval and adult muscles, including the indirect flight muscles. This result suggests that glutamate is indeed the primary neurotransmitter at all adult NMJs, consistent with extensive evidence that *Drosophila* motor neurons release glutamate (i.e., they express the vesicular glutamate transporter, vGlut (Baek and Mann, 2009; Brierley et al., 2012; Daniels et al., 2008)). Thus, some adult muscles may use an alternative glutamate receptor composition that has yet to be characterized. Functional glutamate receptors lacking GluRIIC have not been previously described in vivo, although heterologous reconstitution suggests they are possible. Notably, Han et al. (2015) found that GluRIIA and GluRIIE alone were sufficient to reconstitute functional receptors, without GluRIIC or GluRIID. Because GluRIIA and GluRIIB are thought to occupy the same position within the tetramer, this raises the possibility that GluRIIB-containing complexes lacking GluRIIC — such as the GluRIIB/IID/IIE combination implied by our data in the tibia extensor — could likewise be functional. Neto-β, which is required for receptor gating and synaptic clustering (Han et al., 2024; Kim et al., 2012), is expressed at all adult NMJs we examined (Figures 2D, 3B), so the auxiliary subunit requirement for such complexes would be satisfied. Direct tests of these subunit combinations by heterologous reconstitution are an important next step.

One possibility is that adult muscles express novel or uncharacterized glutamate receptor subunits from among the 16 iGluR genes encoded in the *Drosophila* genome, several of which have not been functionally characterized. A candidate that stands out is *Clumsy*. Our results showed that *Clumsy* is broadly expressed throughout the adult fly, including in muscles controlling the legs, wings, halteres, proboscis, and neck (**Figure s10**). In comparison, we did not detect *Clumsy* in larval muscle. *Clumsy* was originally identified as a mutant with impaired adult locomotion (Berger, 2008; Huen et al., 2001), though this work was never published in a peer-reviewed journal. The gene is phylogenetically related to the GluRIIs (Li et al., 2016), but its function is poorly understood compared to the other subunits expressed in larval muscle. For example, it is not known whether *Clumsy* interacts with *Neto-β.* More work is needed to understand the function of *Clumsy* at the neuromuscular junctions of adult *Drosophila*. Clumsy is not the only candidate. The single-nucleus data indicate that Ukar and Grik are transcribed in adult muscle clusters (**Figures s11, s12**), even though we could not detect expression with the available GAL4 lines. Because indirect flight muscle NMJs were negative for every subunit except Neto-β, these kainate-type receptors merit attention as possible components of adult NMJ receptors. Generating antibodies or endogenously tagged alleles for Clumsy, Ukar, and Grik is a necessary next step toward identifying the receptors that mediate transmission at these synapses.

### Adult abdominal muscles are similar to larvae

Our observation that adult abdominal muscles largely maintain the canonical larval GluRII receptor composition provides important developmental context for understanding receptor diversity. Adult abdominal muscles derive directly from larval body wall muscles through remodeling during metamorphosis, rather than through complete *de novo* muscle formation (Currie and Bate, 1991; Kimura and Truman, 1990). This developmental continuity is reflected in the persistence of the larval receptors, suggesting that the molecular identity established during larval stages is maintained through metamorphosis into muscles that mediate slower abdominal movements (e.g., abdominal pumping during flight (Lehmann and Heymann, 2005)).

In contrast, adult leg and flight muscles arise through *de novo* myogenesis during pupal development (Gunage et al., 2017), providing an opportunity for the developmental program to specify alternative receptor compositions tailored to the biomechanical demands of adult locomotion. The correlation between developmental origin and receptor composition suggests that muscle lineage may contribute to determining synaptic molecular architecture, with metamorphically-remodeled muscles maintaining larval characteristics and newly formed muscles adopting novel molecular identities.

### Extrasynaptic glutamate-gated chloride channels in adult muscle

We identified robust expression of GluClα in adult leg and flight muscles, where the channels localize extrasynaptically along muscle fiber edges rather than at neuromuscular junctions. This finding provides molecular support for classical electrophysiological studies in locusts that described glutamate-gated hyperpolarizing responses (H-receptors) in leg muscles that were functionally and pharmacologically distinct from excitatory D-receptors at the NMJ (Cull-Candy, 1976; Dudel et al., 1989; Lea and Usherwood, 1973). These studies demonstrated that locust leg muscles exhibit two types of glutamate responses: fast depolarizing currents with 1-2 ms kinetics mediated by cation channels at synaptic sites, and slower hyperpolarizing currents with 20-50 ms kinetics mediated by chloride channels distributed extrasynaptically across the muscle surface. Similar observations were made in other invertebrate preparations, including yellow mealworm beetle coxa muscles (Saito and Kawai, 1985) and lobster gastric muscles (Lingle and Marder, 1981), suggesting that extrasynaptic inhibitory glutamate receptors may be a widespread feature of invertebrate muscle physiology. Our identification of GluClα in adult *Drosophila* muscles extends these findings to a genetically tractable organism and raises important questions about the functional role of extrasynaptic inhibitory receptors in motor control.

We note that this arrangement differs from circuits in which excitatory and inhibitory transmission are targeted to different muscles to produce rapid directional control, as in the Mauthner cell escape circuit of larval zebrafish. While most arthropods possess GABAergic motor neurons that directly inhibit leg muscles, holometabolous insects such as *Drosophila* do not (Witten and Truman, 1998). And because GluClα is extrasynaptic rather than apposed to active zones, it is unlikely to support fast, target-specific inhibitory transmission.

We hypothesize that ambient glutamate in the hemolymph, released either from synaptic spillover or from non-neuronal sources, could activate these channels to modulate muscle excitability and tone. By providing tonic hyperpolarization, extrasynaptic GluClα could increase the threshold for muscle activation, potentially preventing inappropriate or excessive contraction. GluClα could also function as a negative feedback controller, in the sense that excess glutamate built up during periods of intense motor activity could activate GluClα to reduce muscle excitability. Ambient glutamate levels in the hemolymph of *Drosophila* larvae are quite high (1-3 mM (Augustin et al., 2007)), while GluClα is fully saturated in oocytes at a glutamate concentration of ∼100 μM (Kane et al., 2000). This feedback mechanism would be particularly important in adult muscles that must sustain precise force control during complex, energetically demanding behaviors like flight and walking.

### Comparison to other animals

Our finding of muscle-specific and developmental regulation of postsynaptic receptor composition parallels glutamatergic synapse heterogeneity in the mammalian brain (Wichmann and Kuner, 2021) and neuromuscular system (Sanes and Lichtman, 1999). During mammalian development, fetal acetylcholine receptors containing the γ subunit are replaced by adult receptors containing the ε subunit at the neuromuscular junction, resulting in altered conductance and channel kinetics (Cetin et al., 2020). Similarly, the distribution of acetylcholine receptors shifts from diffuse expression across the muscle surface to tight clustering at synaptic sites (Sanes and Lichtman, 1999).

While our results reveal that *Drosophila* muscles express extrasynaptic GluClα, mature vertebrate skeletal muscle fibers are not known to express inhibitory neurotransmitter receptors. Instead, they rely on the voltage-gated chloride channel ClC-1 to stabilize the resting membrane potential and prevent hyperexcitability (Pedersen et al., 2016). In vertebrate muscle, extrajunctional acetylcholine receptors are upregulated after injury (Hartzell and Fambrough, 1972), which is thought to increase muscle excitability and facilitate motor neuron reinnervation. Excitatory glutamate receptors are also upregulated after injury in locust leg muscles, while the inhibitory receptors remain unchanged (Cull-Candy, 1975).

### Muscle-specific receptor tuning correlates with biomechanical demands

The heterogeneity in glutamate receptor expression we observed across different muscle types may reflect adaptation to specific biomechanical requirements. Larval body wall muscles mediate slow, rhythmic peristaltic waves against a hydrostatic skeleton, requiring sustained contractions with relatively slow temporal dynamics (0.5-2 Hz, (Heckscher et al., 2012)). In contrast, walking requires precise, graded force control to move jointed limbs against a rigid exoskeleton at higher frequencies (5-20 Hz, (Pratt et al., 2024)). As in other animals, different motor neurons and muscles within the fly leg are specialized for controlling rapid and powerful ballistic movement vs. slow postural stabilization (Azevedo et al., 2020). Different iGluR subunits exhibit a wide range of desensitization kinetics (DiAntonio et al., 1999). Our results raise the possibility that the expression of specific postsynaptic neurotransmitter receptors is tailored to the biomechanical function of different muscles and even fibers within a muscle (e.g., slow vs. fast-twitch).

Fly flight demands even higher frequency muscle contraction. To achieve this, the indirect flight muscles are asynchronous: the motor neurons spike at ∼6 Hz while the stretch-activated muscles contract at >200 Hz (Dickinson and Tu, 1997). In contrast, direct flight muscles at the base of the wing hinge provide fine-tuned steering control through synchronous contraction with motor neuron spiking. While we did not detect canonical GluRs in the indirect flight muscles, the direct flight muscles do faintly express GluRIIA. The differential receptor expression between direct and indirect flight muscles may reflect their distinct functional roles: power production vs. steering may require unique receptor kinetics, conductances, and desensitization properties.

Another major finding from larval NMJ studies is presynaptic homeostasis, wherein reduction of postsynaptic GluRs triggers compensatory increases in presynaptic neurotransmitter release. Our finding that canonical postsynaptic GluRII receptors were not detected in adult leg and flight muscles raises questions about whether and how homeostatic mechanisms operate at these specialized synapses. Do muscles without detectable canonical iGluR receptors retain the capacity for presynaptic homeostatic potentiation? If so, what postsynaptic sensors and retrograde signals mediate this plasticity?

Alternatively, do highly specialized muscles trade plasticity for precision, relying on fixed synaptic properties that are established during development and remain relatively stable throughout adult life?

### Limitations

Our study employed multiple complementary approaches to characterize receptor expression patterns. The convergence of results across GAL4 reporter expression, antibody staining, endogenous tagging, and single-cell RNA-seq data supports our conclusions about differences between larval and adult muscles. However, several technical limitations should be noted. GAL4 reporters may not capture all aspects of endogenous gene regulation. Antibody specificity, while validated in larval muscles, may differ in the adult. Single-cell RNA-seq provides a snapshot of the transcriptome but does not directly assess protein expression or localization. In the future, electrophysiological recordings from adult NMJs and GluR mutant characterization will be essential to more directly assess synaptic transmission properties and to confirm the functional consequences of the receptor expression patterns we describe.

Inferring the absence of a protein from the absence of signal requires caution, because every method we used is subject to false negatives. Several of our negative results, however, are internally controlled. In the femur, anti-GluRIIB labels NMJs on tibia extensor fibers but not on accessory tibia flexor fibers within the same bisected femur, while anti-GluRIIC shows the reciprocal pattern in the same preparations (Figure 3C-F); the failure to detect a subunit in one muscle therefore cannot be explained by a general failure of that antibody in adult tissue. Within the tibia extensor, anti-GluRIIB labels proximal but not distal fibers, an internal control at the scale of a single muscle. In indirect flight muscle, where we detected none of the GluRII subunits, Brp, Neto-β::sfGFP, and GluClα:V5 were all readily detected in the same preparations, demonstrating that reagents access this tissue and that NMJs are correctly identified. These internal controls do not exclude receptor expression below our detection threshold, and we therefore describe our results throughout as “not detected” rather than “absent.” They do, however, indicate that the muscle-specific differences we report reflect biological variation rather than technical failure.

We did observe some discrepancies across the different experimental approaches. For GAL4 reporter lines, we used T2A gene trap lines where possible (for 15 of the 16 GluR genes) to determine receptor expression. However, some of these reporters were extremely weak and may have resulted in false negatives. For example, we did not detect Grik-GAL4 or Ukar-GAL4 expression in any tissue **(Figure 2C)**, despite RNA-seq evidence that these are expressed robustly in adult muscles **(Figures s11, s12)**. We do not have antibodies or protein reporters for these genes to help resolve this. Another discrepancy was with GluRIIE. In this case, antibody staining and RNA-seq did not detect GluRIIE in flight muscles **(Figures s8, s11)**, while the GAL4>GFP reporter showed extremely faint signal in the flight muscles. Because of the marginal signal and the discrepancy with the other methods, we believe this could likely be a false positive. The table in Figure 2 does not effectively capture the nuance of this otherwise confusing result. Overall, however, we found that results from different approaches generally agreed and revealed intriguing differences between different muscle types.

### Conclusions

Our findings highlight the importance of examining neuromuscular function in appropriate developmental and anatomical context. More broadly, our results emphasize that developmental stage and tissue identity shape synaptic molecular architecture. As the field moves toward comprehensive, multi-scale models of motor control that integrate connectomics, biomechanics, and neural dynamics, it will be important to define the molecular and functional properties of synaptic transmission for different muscles. Understanding neuromuscular diversity and the principles that govern it will be essential for building accurate models of motor control and for understanding how the nervous system adapts its output to match the mechanical properties of the effectors it must control.

## Acknowledgements and Support

We thank members of the Tuthill and Brunton Labs for technical assistance and feedback on the manuscript, especially Tony Azevedo for his help with understanding leg musculature. We thank Himanshu Gupta for sharing unpublished results and helpful discussions about leg motor neurons. We acknowledge the UW Keck Microscopy Center and its NIH S10 funding (S10 OD016240). We acknowledge the Developmental Studies Hybridoma Bank (Iowa, USA) for antibodies used in this study, and the Bloomington Drosophila Stock Center for fly stocks (NIH P40OD018537). Financial support was provided by NIH grant R01NS128785, a Searle Scholar Award, a Klingenstein-Simons Fellowship, a Pew Biomedical Scholar Award, a McKnight Scholar Award, a Sloan Research Fellowship, the New York Stem Cell Foundation, and a UW Innovation Award to J.C.T., and NIH grant R01NS126654 and NSF grant IOS-2417451 to D.D. J.C.T is a New York Stem Cell Foundation – Robertson Investigator.

## Contributions

AS and JCT conceived the study. AS collected data, performed analyses, and drafted figures. CQ, YX, and DD generated transgenic fly stocks and antibodies. AS, CQ, DD, and JCT interpreted the results. AS and JCT wrote the paper with input from CQ and DD.

## Materials and Methods

### Fly husbandry

Flies were raised at 25°C on standard fly food with cornmeal, agar, and molasses. We dissected animals for analysis at the third instar larvae stage and at 2-4 days after adult eclosion. In cases where we failed to detect any GAL4 signal (e.g., GluRIID-GAL4), we additionally imaged very young and very old flies (0-2 hours old, and 20 day-old). It did not change the results.

### Sample preparation for confocal imaging of GAL4/UAS in hemisected adult flies

For imaging GFP expression in hemisected adult flies, we fixed whole female flies in fixative (4% paraformaldehyde in PBS). Next, flies were one by one transferred to a small drop of Tissue-Tek O.C.T., frozen on a glass slide on dry ice for 15 seconds, sliced along the anterior-posterior axis with a fine razor blade and transferred to a well with fixative for one minute, until the O.C.T. melted and dissolved. The tissue was then washed 3 times in PBS containing 0.2% Triton-X (PBST) over one hour. Tissue was cleared in FocusClear for 20 minutes and mounted in Mount Clear on a slide with four layers of 3M double-stick tape for spacers. Preps were imaged on a Confocal Olympus FV1000.

For imaging of whole larvae, we transferred larvae to a drop of Vectashield on a glass slide with double-stick tape spacers. We froze the larvae in Vectashield for 5 minutes to immobilize them, then covered the larvae with a coverslip, and immediately imaged them on a Leica DMI6000 widefield microscope (UW Keck Imaging Center). Images were processed in FIJI (Schindelin et al., 2012).

### Immunohistochemistry and NMJ imaging

The tissue was fixed in the same manner as above. Femurs were bisected longitudinally in O.C.T. to allow reagent penetration. All fixing and staining steps were at room temperature with gentle nutation. Tissue was put into a blocking solution (5% goat serum, PBS, 0.2% Triton-X) for 1 hour, then primary antibody solution for 24 hours. Tissue was washed three times in PBST and incubated in blocking and secondary antibody solution (see concentrations in the table below). Finally, tissue was incubated overnight in Alexa Fluor 647-nm Phalloidin in a PBS solution with the following reagents to improve tissue penetrance: 1% triton X-100, 0.5% DMSO, 0.05 mg/ml escin, and 3% normal goat serum. Tissue was washed three times with PBST over three hours, once in PBS for one hour, and mounted on a slide in Vectashield. NMJ images were collected on a Leica SP8 confocal (UW Keck Imaging Center). Images were processed in FIJI (Schindelin et al., 2012).

### Quantification of GAL4/UAS and IHC expression for tables in Figures 2 and s14

For characterizing GAL4/UAS expression in whole larva and whole flies, we fixed and mounted a minimum of six animals per genotype. For each tissue type, we collected two 20x confocal stacks from two representative flies per genotype. (In no cases did expression patterns vary significantly; see **Figure s9**.) We examined the confocal stacks in FIJI and quantified the expression intensity and fraction of muscles qualitatively: three levels for intensity (undetected, weak, strong) and five levels for approximate fraction of muscles (0, 25%, 50%, 75%, or 100%).

For quantifying protein expression at neuromuscular junctions, we stained and mounted a minimum of six animals per genotype. We used a low power objective to identify general muscle regions (e.g. coxa, indirect flight muscle). We then zoomed in with a 60x microscope objective to find and collect images where an NMJ was in good view, with strong control stain (Brp) and in a focal plane close to the microscope objective for best resolution. For each staining condition and tissue type, we collected a minimum of four representative image stacks. As above, expression intensity and approximate fraction of NMJ expression was quantified qualitatively in FIJI.

To document muscle-specific GluR receptors within the femur, we mounted and collected confocal 20x stacks of ten femurs per experiment, then used the 60x objective with a 3x digital zoom to collect stacks of NMJs in specific muscle fibers. We analyzed the images qualitatively for receptor expression.

### RNA-seq analysis

We accessed single-cell RNA-seq Fly Cell Atlas datasets through the SCope website (Davie et al., 2018; Li et al., 2022). Tissue types were determined based on Fly Cell Atlas annotation. **Figure s11** shows the 10x Cross-tissue/Muscle system/Stringent dataset. **Figure s12** shows the 10x Leg/Stringent dataset.

### Tagged Neto and GluRIIB proteins

CRISPR/Cas9-mediated homology-directed repair (HDR) was performed by WellGenetics Inc. (Taipei City, 11570, Taiwan) to target the Neto (CG44328) locus to generate endogenously tagged Neto-α and Neto-β isoforms. The gRNA sequences GAACTTGAGGGGATGTACGC and GTCCCATTGACATTTAAGGA were cloned into U6 promoter plasmids to target Neto-α and Neto-β, respectively. For both isoforms, tag cassettes were inserted immediately upstream of the C-terminal stop codon via homology-directed repair using pUC57-Kan donor vectors containing flanking homology arms. The Neto-α donor contained an mScarlet-PBacGFP cassette (mScarlet, 3xGly, 3xFLAG, and a floxed 3xP3-GFP), while the Neto-β donor contained an sfGFP-3xP3-RFP cassette (sfGFP and a floxed 3xP3-RFP). These donors were microinjected into w^1118^ embryos alongside the respective gRNA plasmids and hs-Cas9. Successful F1 transformants were identified by marker expression (3xP3-GFP or 3xP3-RFP) and validated by genomic PCR and sequencing. Subsequently, the selection markers were excised from the confirmed lines using piggyBac transposase.

To generate the endogenously tagged GluRIIB::ALFA allele, a gRNA (TTCGACGGGTTACTCATCGC) targeting the GluRIIB C-terminus was cloned into a pU6 plasmid by GenScript (Piscataway, NJ 08854). A single-stranded DNA (ssDNA) donor template was designed to insert an ALFA tag flanked by a proline linker and homology arms. To prevent Cas9 cleavage of the donor and the re-cleavage of the edited locus, two nucleotide mutations were introduced into the left homology arm to disrupt gRNA binding. The full reverse-complement ssDNA donor sequence was synthesized (TTTTTACTCCCCTACTTTTCAATTCGCCTGGTCTTTTTTTTCTTTTTCGCACTGGAAGCAGGTTCTGTGAGCCGTCGCCTCAGCTCTTCTTCCAGCCTACTTGGACTTGTAATTTGCTCCAAGGATGAGTAACCCGTCGAAGTTGAATCCCGACTTTGGCGATAGGTGTGCAGCGGTTTTCT) by Integrated DNA Technologies (IDT; Coralville, IA 52241). The gRNA plasmid and ssDNA donor were co-injected into vas-Cas9 embryos (BDSC #51324) by Bestgene (Chino Hills, CA 91709). Correct insertions were identified via PCR screening using the following primers: GATTCGGTGGGTGGACTGTTC (forward) and CGTATTTCACTGAATGTTCGCAAGC (reverse). Successful editing was subsequently confirmed by Sanger sequencing and immunostaining.

### GluRIIE antibody

To generate the anti-GluRIIE antibody, the peptide C-SASIYSRSRQSSMSVASVAQESQ was synthesized by Bio-Synthesis, Inc. (Lewisville, TX 75057, USA). This antigen was used for rabbit immunization and subsequent affinity purification of the sera by Cocalico Biologicals (Denver, PA 17517, USA).

### Antibody validation

The anti-GluRIIB, anti-GluRIIC, and anti-GluRIID antibodies used here were generated and validated previously against loss-of-function alleles and RNAi knockdown at the larval NMJ (Goel and Dickman, 2018; Perry et al., 2017). The anti-GluRIIE antibody was generated for this study and tested against a CRISPR allele that truncates the GluRIIE C-terminus, removing the antigenic sequence. Signal at larval NMJs was strongly reduced but not eliminated in this mutant, with a faint residual signal that we attribute to cross-reactivity with GluRIID, whose C-terminal sequence is similar. We note that this cross-reactivity makes the antibody conservative for the negative results reported here: in tissues where we detected no signal, such as the indirect flight muscles, neither GluRIIE nor GluRIID was detected by any approach. Conversely, the faint GluRIIE signal in adult abdominal muscle (Figure s8) should be interpreted with the possibility of GluRIID cross-reactivity in mind. For all experiments in adult tissue, larval preparations were stained in parallel using the same antibody dilutions, incubation times, and imaging settings, so that larval NMJs served as a positive control for each staining run.

### Table of Fly Stocks

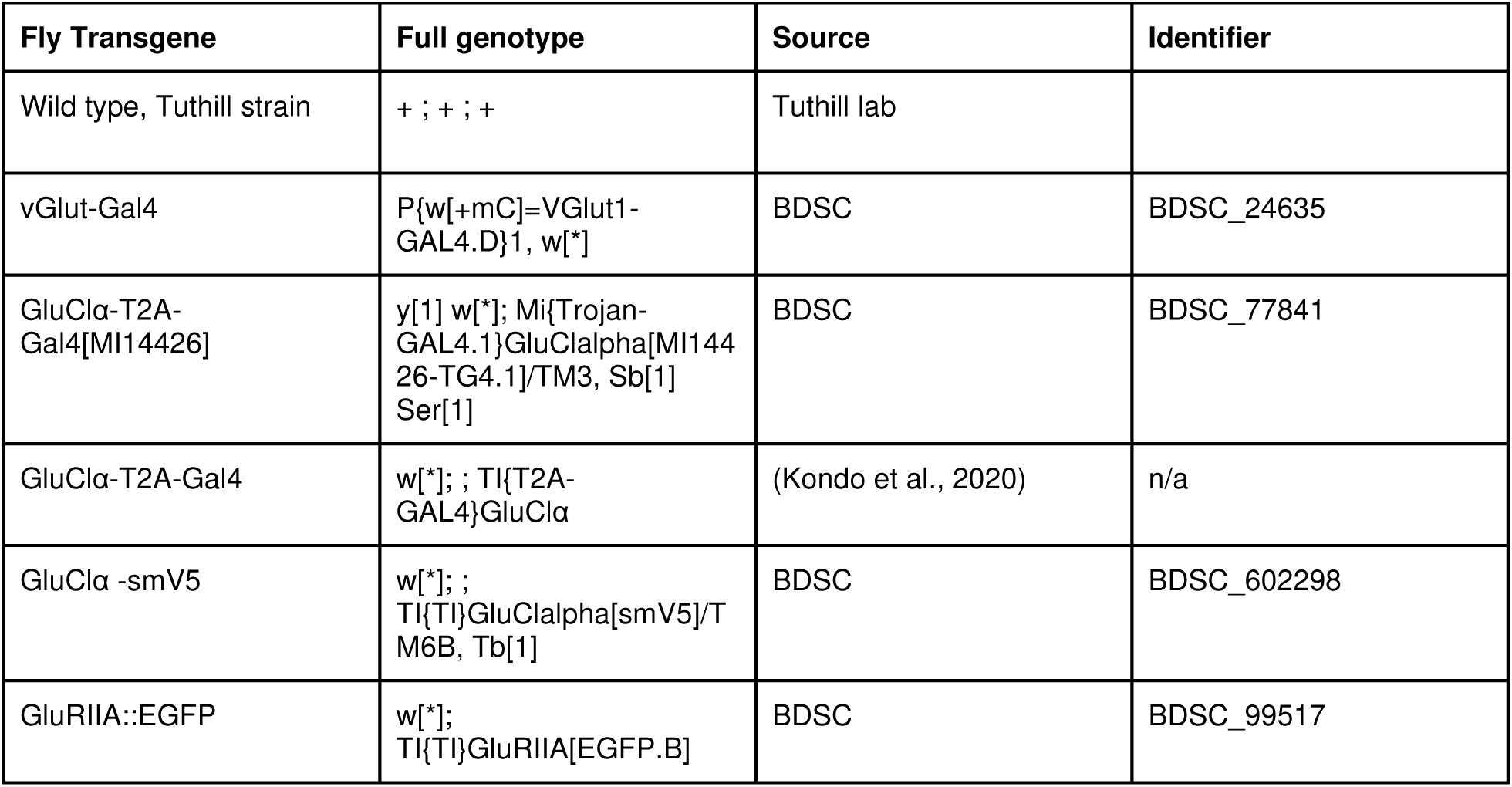
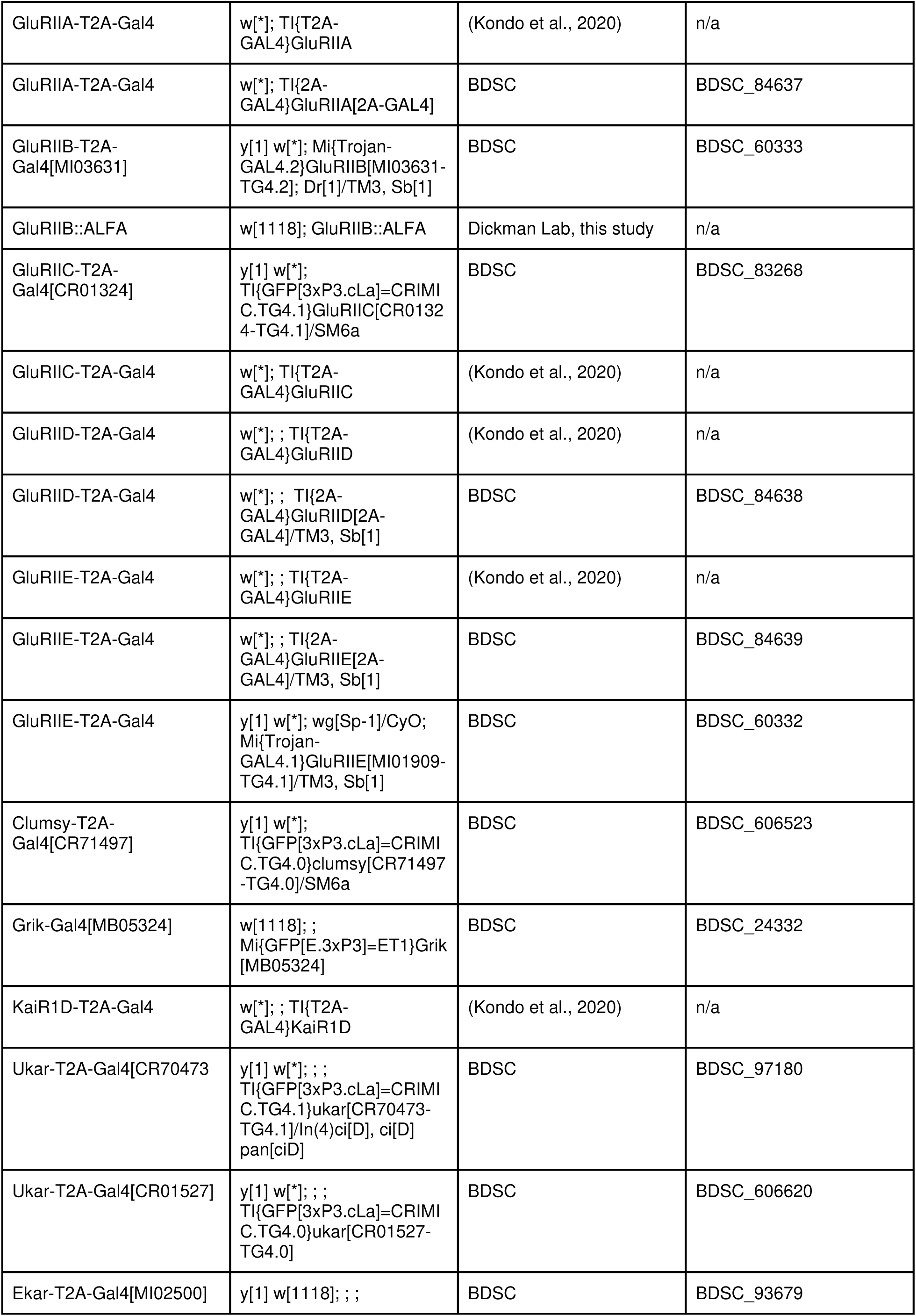
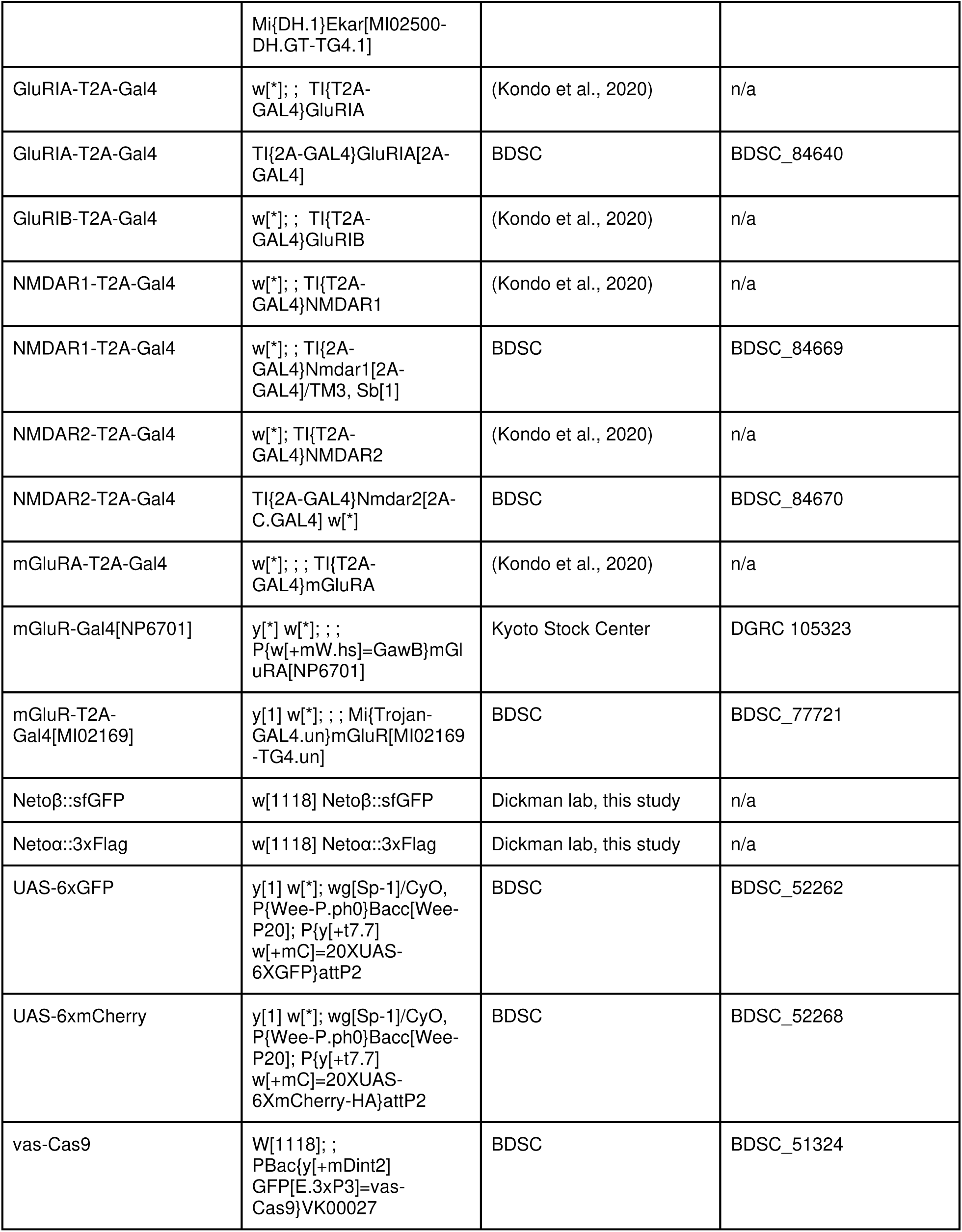

### Key Resources Table

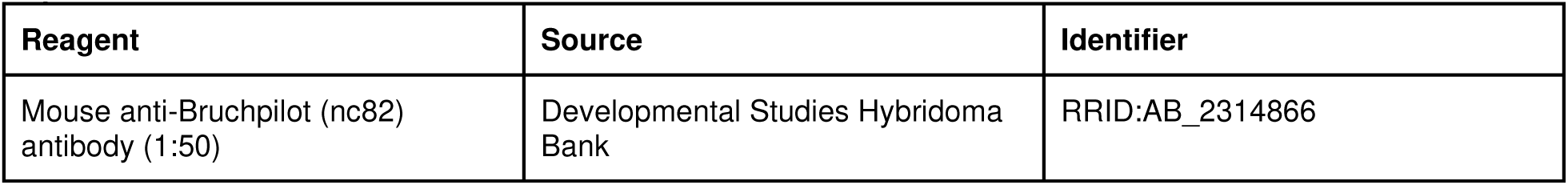
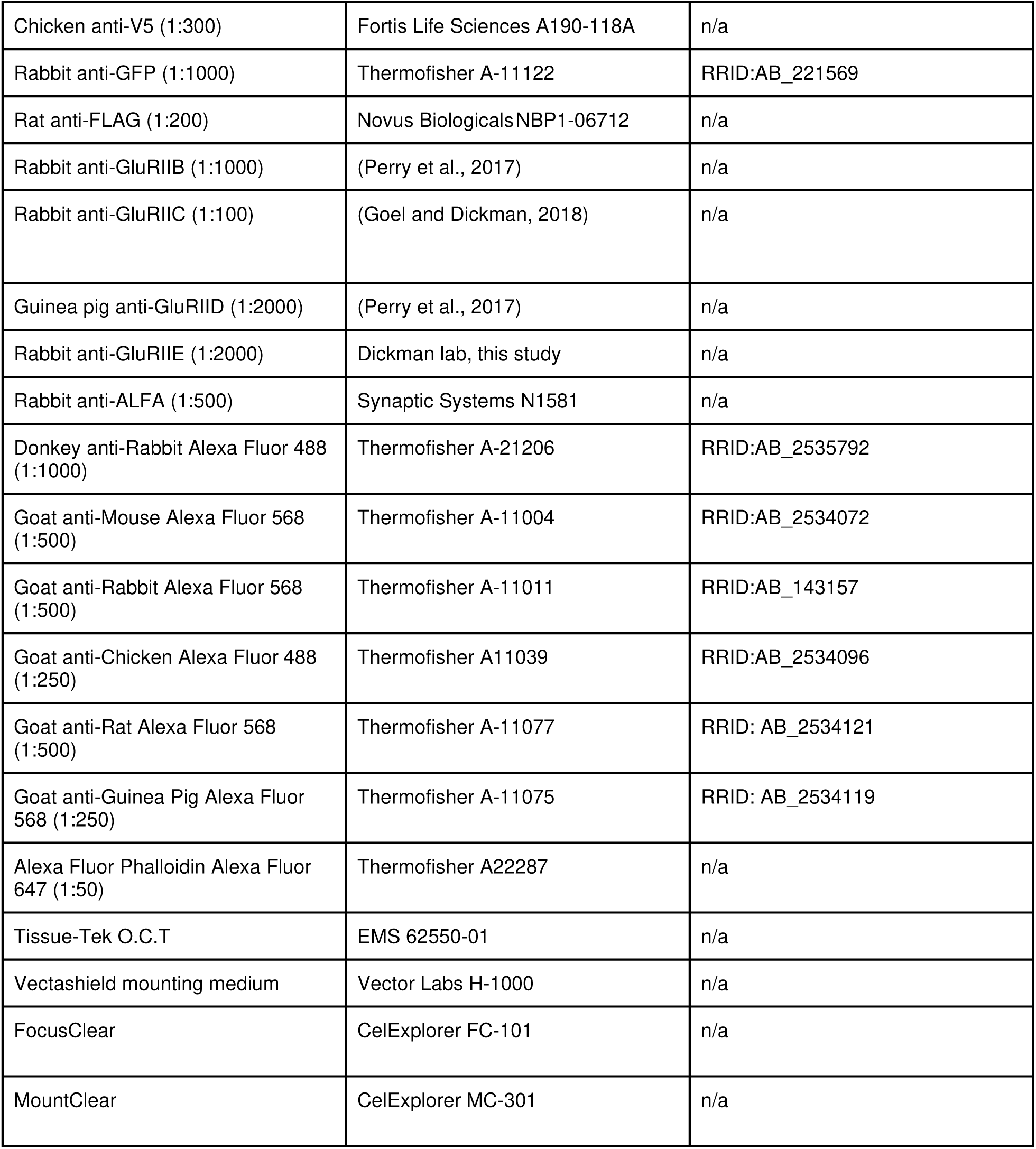

| Reagent | Source | Identifier |
| --- | --- | --- |
| Mouse anti-Bruchpilot (nc82) antibody (1:50) | Developmental Studies Hybridoma Bank | RRID:AB_2314866 |
| Chicken anti-V5 (1:300) | Fortis Life Sciences A190-118A | n/a |
| Rabbit anti-GFP (1:1000) | Thermofisher A-11122 | RRID:AB_221569 |
| Rat anti-FLAG (1:200) | Novus Biologicals NBP1-06712 | n/a |
| Rabbit anti-GluRIIB (1:1000) | (Perry et al., 2017) | n/a |
| Rabbit anti-GluRIIC (1:100) | (Goel and Dickman, 2018) | n/a |
| Guinea pig anti-GluRIID (1:2000) | (Perry et al., 2017) | n/a |
| Rabbit anti-GluRIIE (1:2000) | Dickman lab, this study | n/a |
| Rabbit anti-ALFA (1:500) | Synaptic Systems N1581 | n/a |
| Donkey anti-Rabbit Alexa Fluor 488 (1:1000) | Thermofisher A-21206 | RRID:AB_2535792 |
| Goat anti-Mouse Alexa Fluor 568 (1:500) | Thermofisher A-11004 | RRID:AB_2534072 |
| Goat anti-Rabbit Alexa Fluor 568 (1:500) | Thermofisher A-11011 | RRID:AB_143157 |
| Goat anti-Chicken Alexa Fluor 488 (1:250) | Thermofisher A11039 | RRID:AB_2534096 |
| Goat anti-Rat Alexa Fluor 568 (1:500) | Thermofisher A-11077 | RRID: AB_2534121 |
| Goat anti-Guinea Pig Alexa Fluor 568 (1:250) | Thermofisher A-11075 | RRID: AB_2534119 |
| Alexa Fluor Phalloidin Alexa Fluor 647 (1:50) | Thermofisher A22287 | n/a |
| Tissue-Tek O.C.T | EMS 62550-01 | n/a |
| Vectashield mounting medium | Vector Labs H-1000 | n/a |
| FocusClear | CelExplorer FC-101 | n/a |
| MountClear | CelExplorer MC-301 | n/a |

### Table of Genotypes

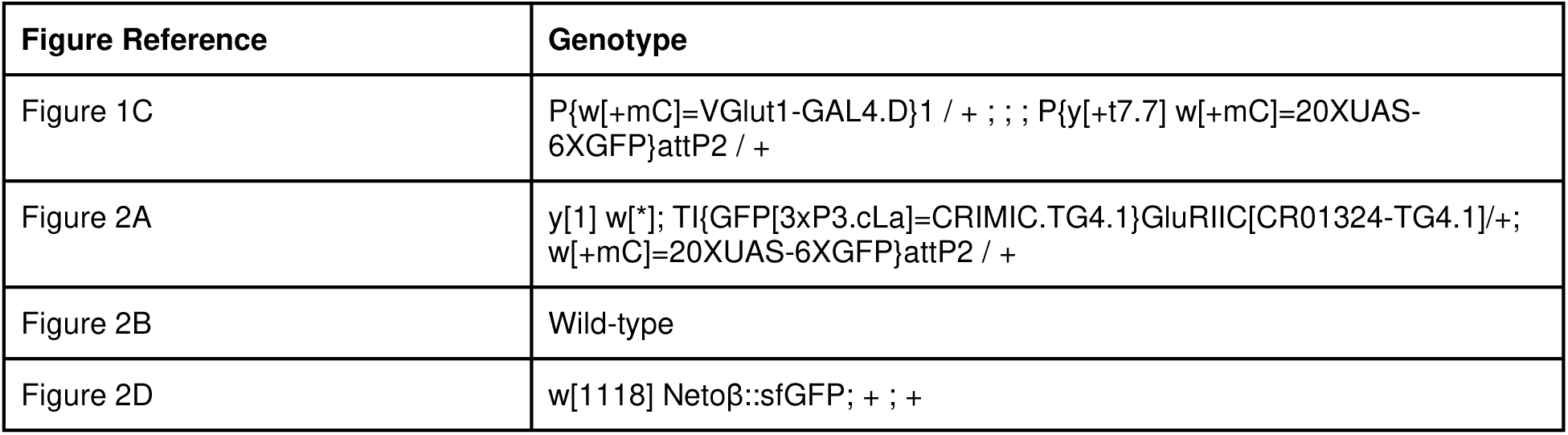
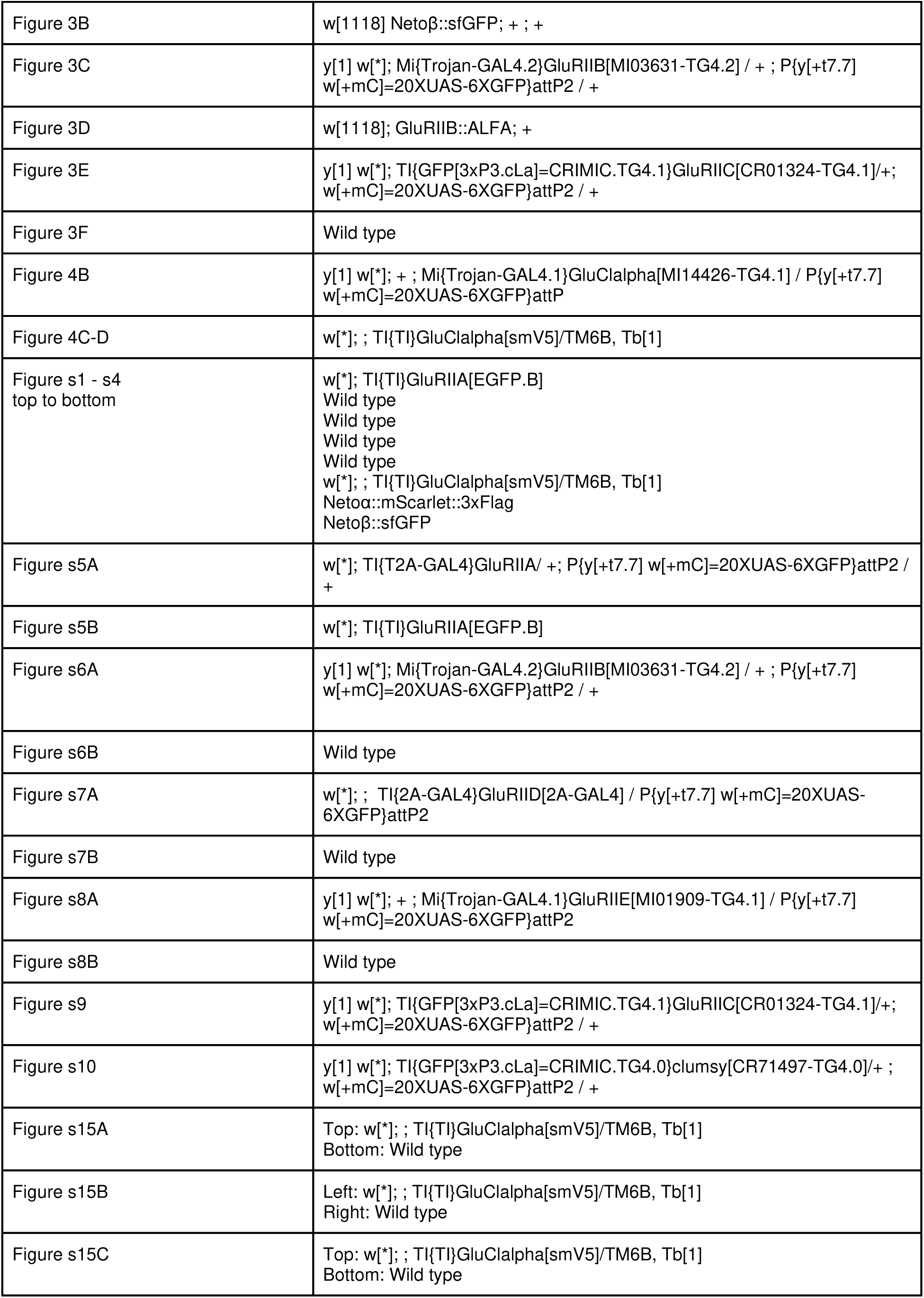

| Figure Reference | Genotype |
| --- | --- |
| Figure 1C | P{w[+mC]=VGlut1-GAL4.D}1 / + ; ; ; P{y[+t7.7] w[+mC]=20XUAS-6XGFP}attP2 / + |
| Figure 2A | y[1] w[*]; TI{GFP[3xP3.cLa]=CRIMIC.TG4.1}GluRIIC[CR01324-TG4.1]/+; w[+mC]=20XUAS-6XGFP}attP2 / + |
| Figure 2B | Wild-type |
| Figure 2D | w[1118] Netoβ::sfGFP; + ; + |
| Figure 3B | w[1118] Netoβ::sfGFP; + ; + |
| Figure 3C | y[1] w[*]; Mi{Trojan-GAL4.2}GluRIIB[Mi03631-TG4.2] / + ; P{y[+t7.7] w[+mC]=20XUAS-6XGFP}attP2 / + |
| Figure 3D | w[1118]; GluRIIB::ALFA; + |
| Figure 3E | y[1] w[*]; TI{GFP[3xP3.cLa]=CRIMIC.TG4.1}GluRIIC[CR01324-TG4.1]/+; w[+mC]=20XUAS-6XGFP}attP2 / + |
| Figure 3F | Wild type |
| Figure 4B | y[1] w[*]; + ; Mi{Trojan-GAL4.1}GluClalpha[Mi14426-TG4.1] / P{y[+t7.7] w[+mC]=20XUAS-6XGFP}attP |
| Figure 4C-D | w[*]; ; TI{TI}GluClalpha[smV5]/TM6B, Tb[1] |
| Figure s1 - s4<br>top to bottom | w[*]; TI{TI}GluRIIA[EGFP.B]<br>Wild type<br>Wild type<br>Wild type<br>Wild type<br>w[*]; ; TI{TI}GluClalpha[smV5]/TM6B, Tb[1]<br>Netoα::mScarlet::3xFlag<br>Netoβ::sfGFP |
| Figure s5A | w[*]; TI{T2A-GAL4}GluRIIA/ +; P{y[+t7.7] w[+mC]=20XUAS-6XGFP}attP2 / + |
| Figure s5B | w[*]; TI{TI}GluRIIA[EGFP.B] |
| Figure s6A | y[1] w[*]; Mi{Trojan-GAL4.2}GluRIIB[Mi03631-TG4.2] / + ; P{y[+t7.7] w[+mC]=20XUAS-6XGFP}attP2 / + |
| Figure s6B | Wild type |
| Figure s7A | w[*]; ; TI{2A-GAL4}GluRIID[2A-GAL4] / P{y[+t7.7] w[+mC]=20XUAS-6XGFP}attP2 |
| Figure s7B | Wild type |
| Figure s8A | y[1] w[*]; + ; Mi{Trojan-GAL4.1}GluRIIE[Mi01909-TG4.1] / P{y[+t7.7] w[+mC]=20XUAS-6XGFP}attP2 |
| Figure s8B | Wild type |
| Figure s9 | y[1] w[*]; TI{GFP[3xP3.cLa]=CRIMIC.TG4.1}GluRIIC[CR01324-TG4.1]/+; w[+mC]=20XUAS-6XGFP}attP2 / + |
| Figure s10 | y[1] w[*]; TI{GFP[3xP3.cLa]=CRIMIC.TG4.0}clumsy[CR71497-TG4.0]/+ ; w[+mC]=20XUAS-6XGFP}attP2 / + |
| Figure s15A | Top: w[*]; ; TI{TI}GluClalpha[smV5]/TM6B, Tb[1]<br>Bottom: Wild type |
| Figure s15B | Left: w[*]; ; TI{TI}GluClalpha[smV5]/TM6B, Tb[1]<br>Right: Wild type |
| Figure s15C | Top: w[*]; ; TI{TI}GluClalpha[smV5]/TM6B, Tb[1]<br>Bottom: Wild type |

