## supplemental figures for "Glutamate receptor composition at *Drosophila* neuromuscular junctions depends on developmental stage and muscle identity"

Sustar et al., 2026. Supplemental Figures.

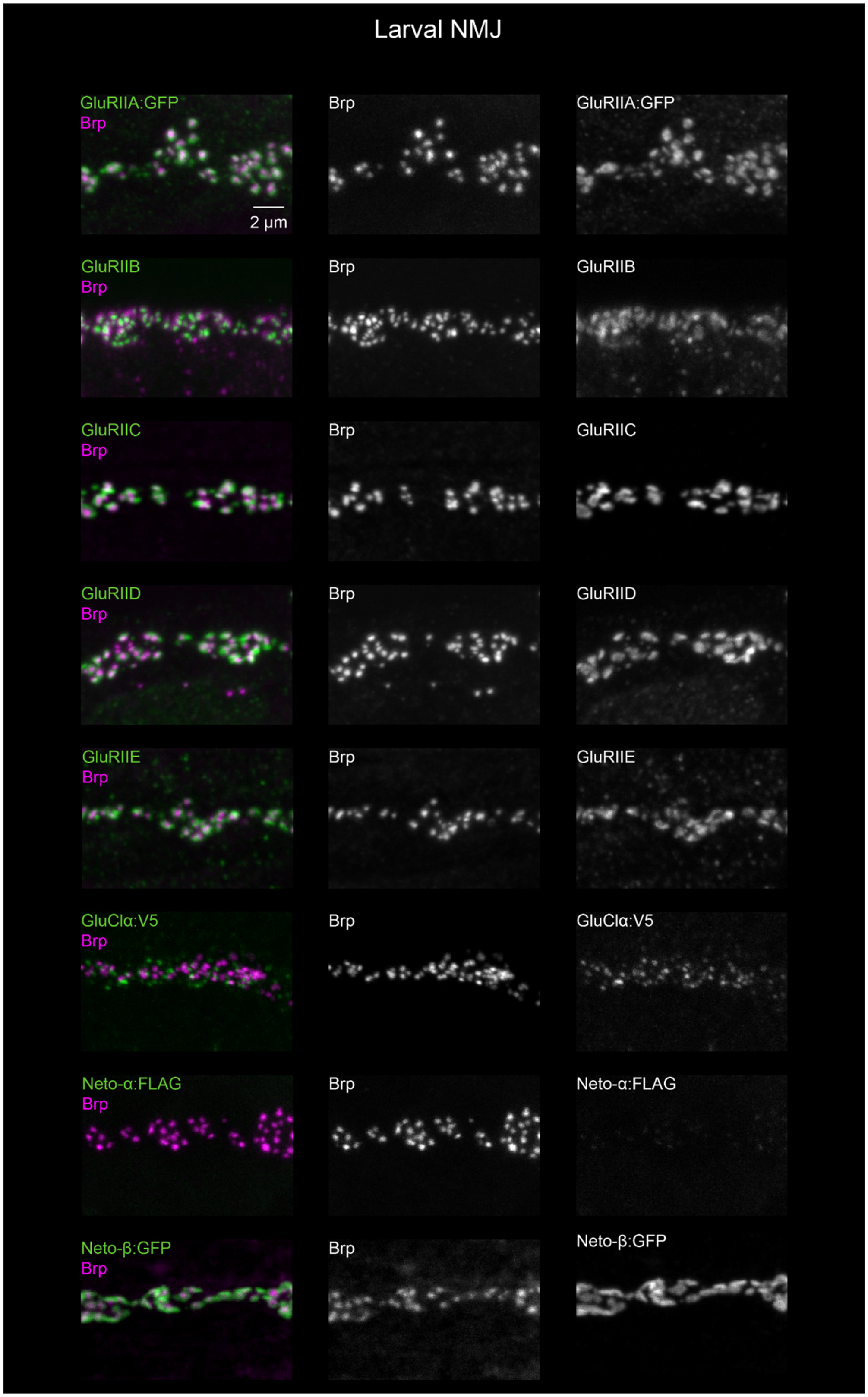


**Figure s1.** NMJ staining of larval body wall muscle.

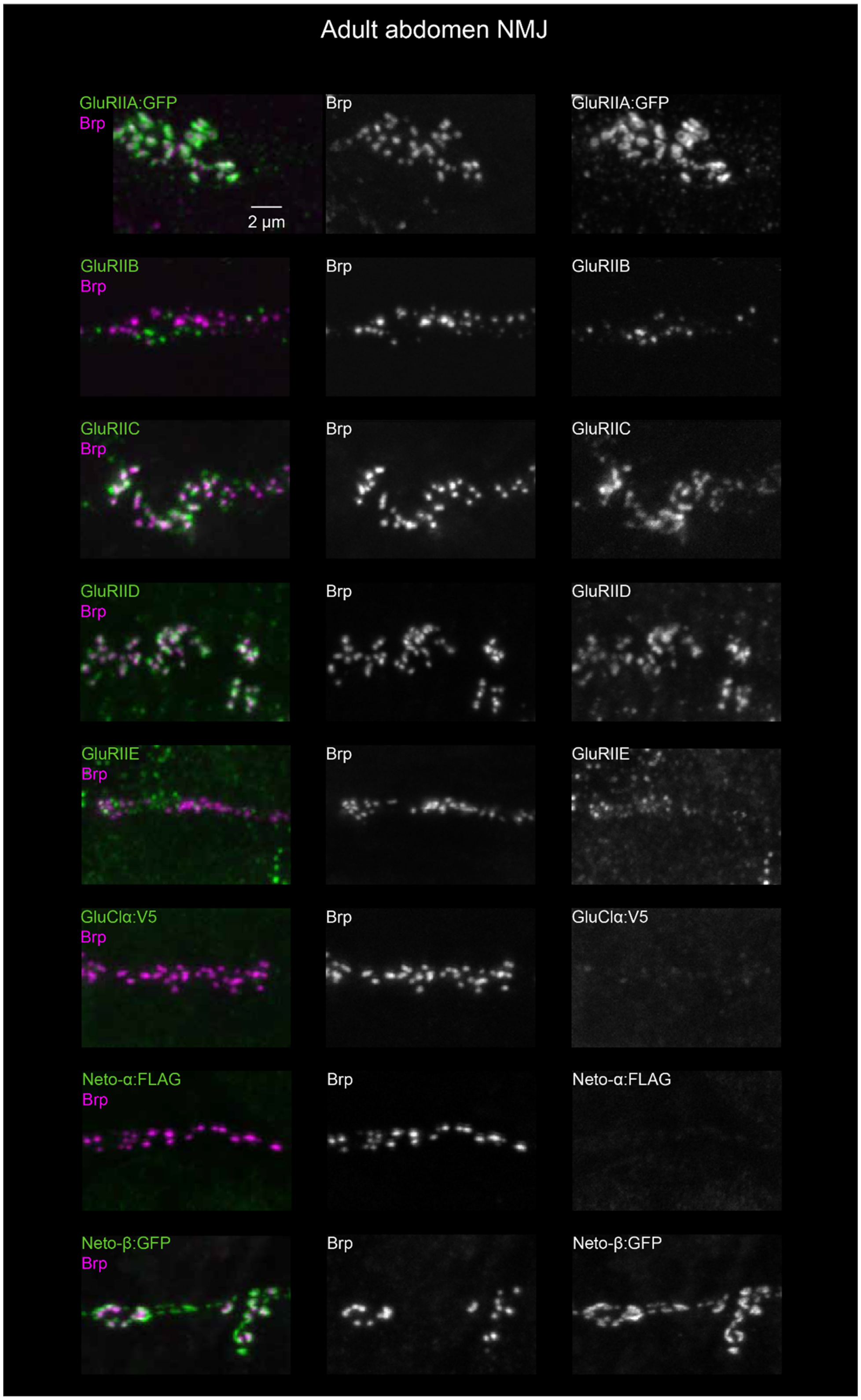


**Figure s2.** NMJ staining of adult abdomen muscle.

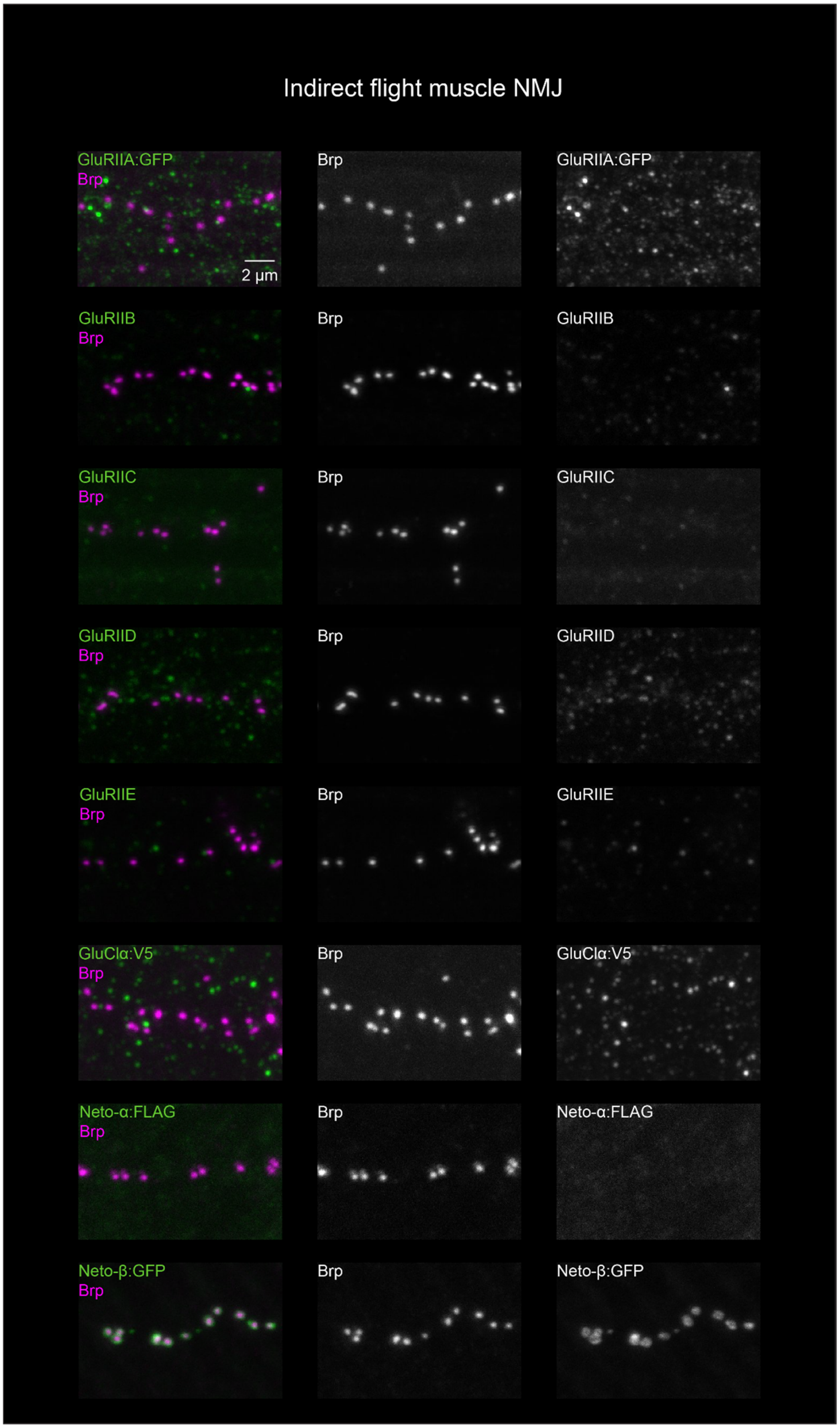


**Figure s3.** NMJ staining of adult indirect flight muscle.

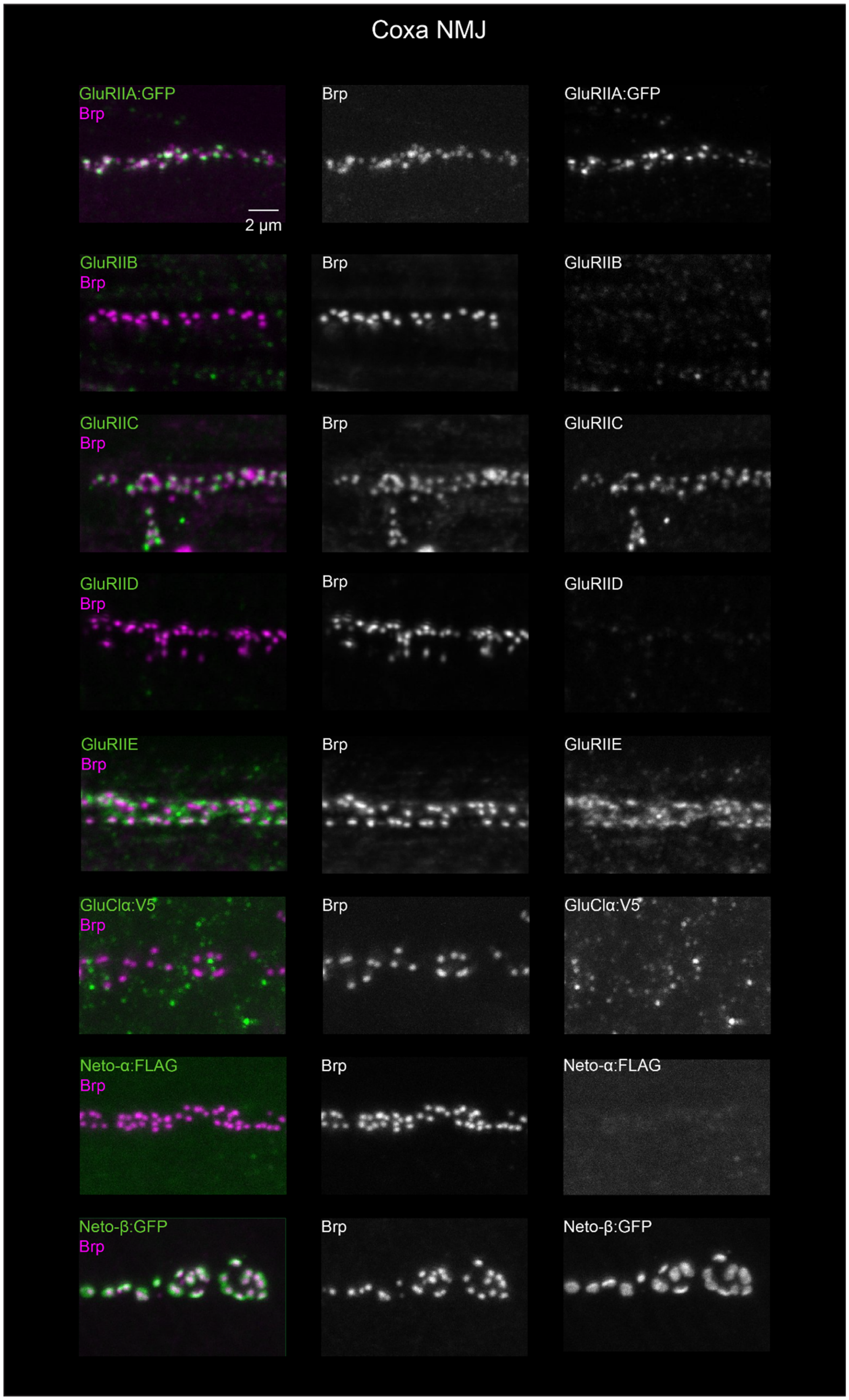


**Figure s4.** NMJ staining of adult leg muscle (coxa).

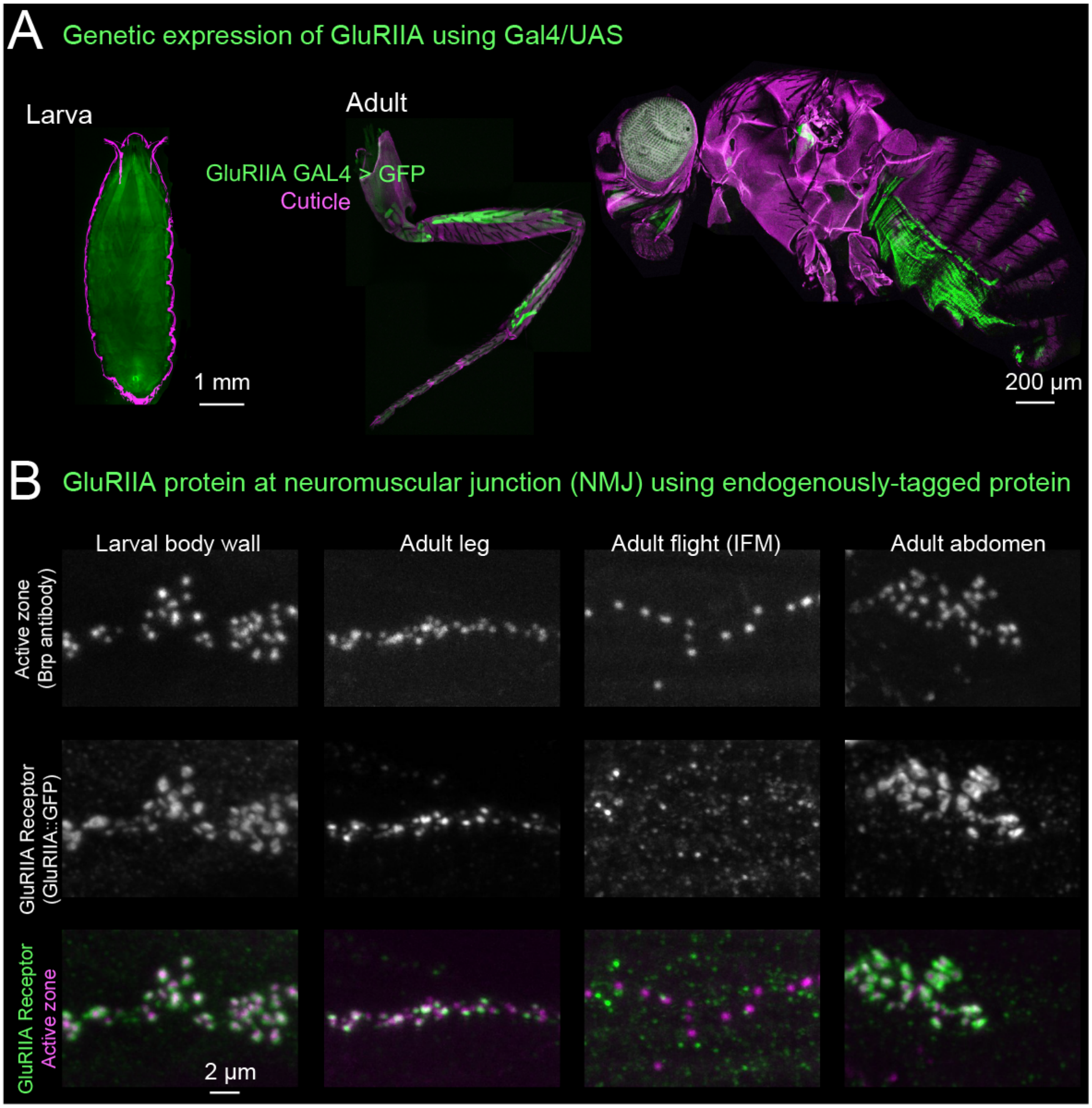


**Figure s5.** (A) Representative confocal images showing GFP expression driven by a GAL4 reporter line (GluRIIA-GAL4) in larval and adult muscles. (B) Representative images showing antibody staining of GluRIIA::GFP protein (green) at larval and adult neuromuscular junctions (magenta).

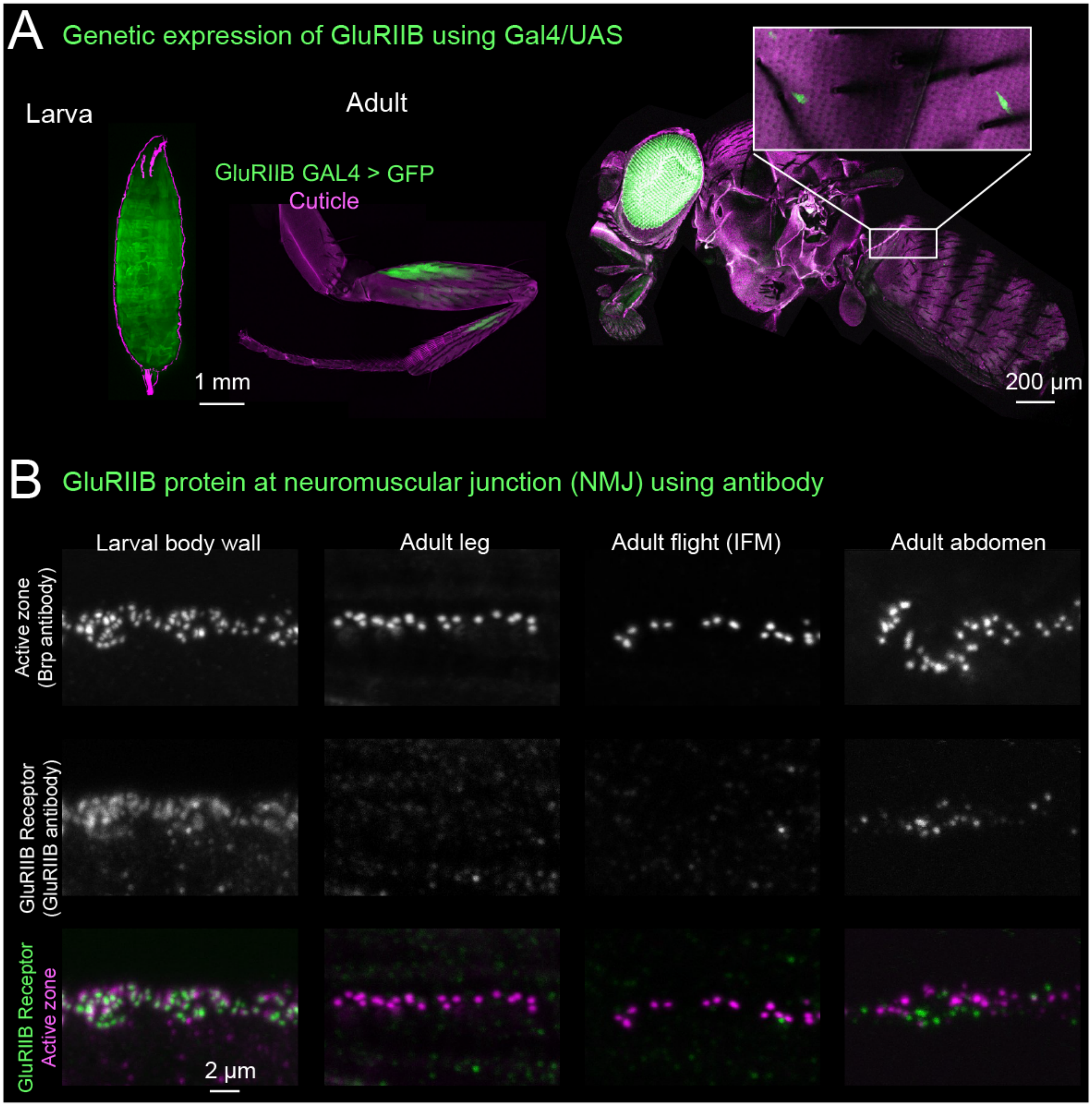


**Figure s6.** (A) Representative confocal images showing GFP expression driven by a GAL4 reporter line (GluRIIB-GAL4) in larval and adult muscles. (B) Representative images showing antibody staining of GluRIIB protein (green) at larval and adult neuromuscular junctions (magenta).

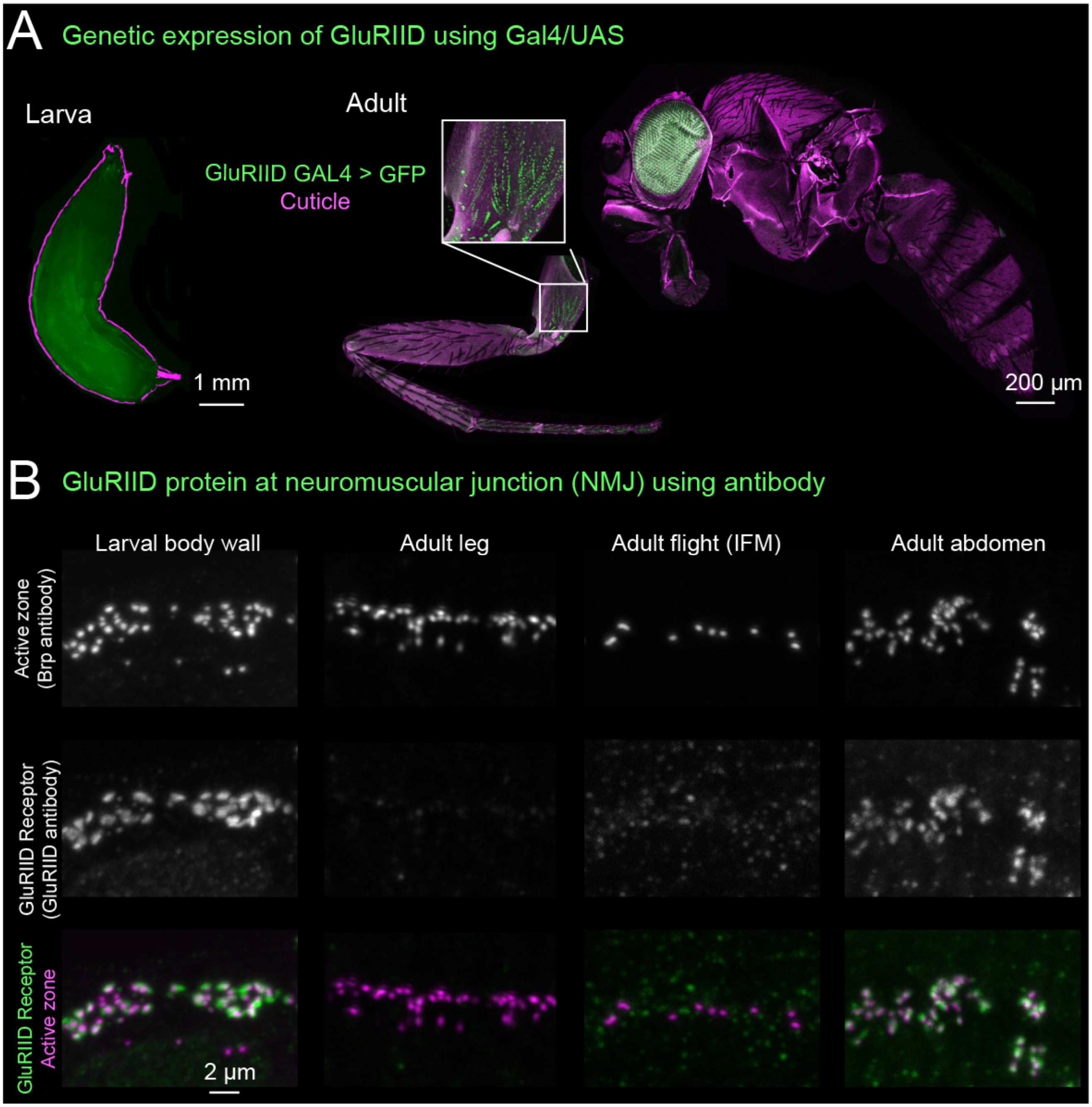


**Figure s7.** (A) Representative confocal images showing GFP expression driven by a GAL4 reporter line (GluRIID-GAL4, green) in larval and adult muscles. (B) Representative images showing antibody staining of GluRIID protein (green) at larval and adult neuromuscular junctions (magenta).

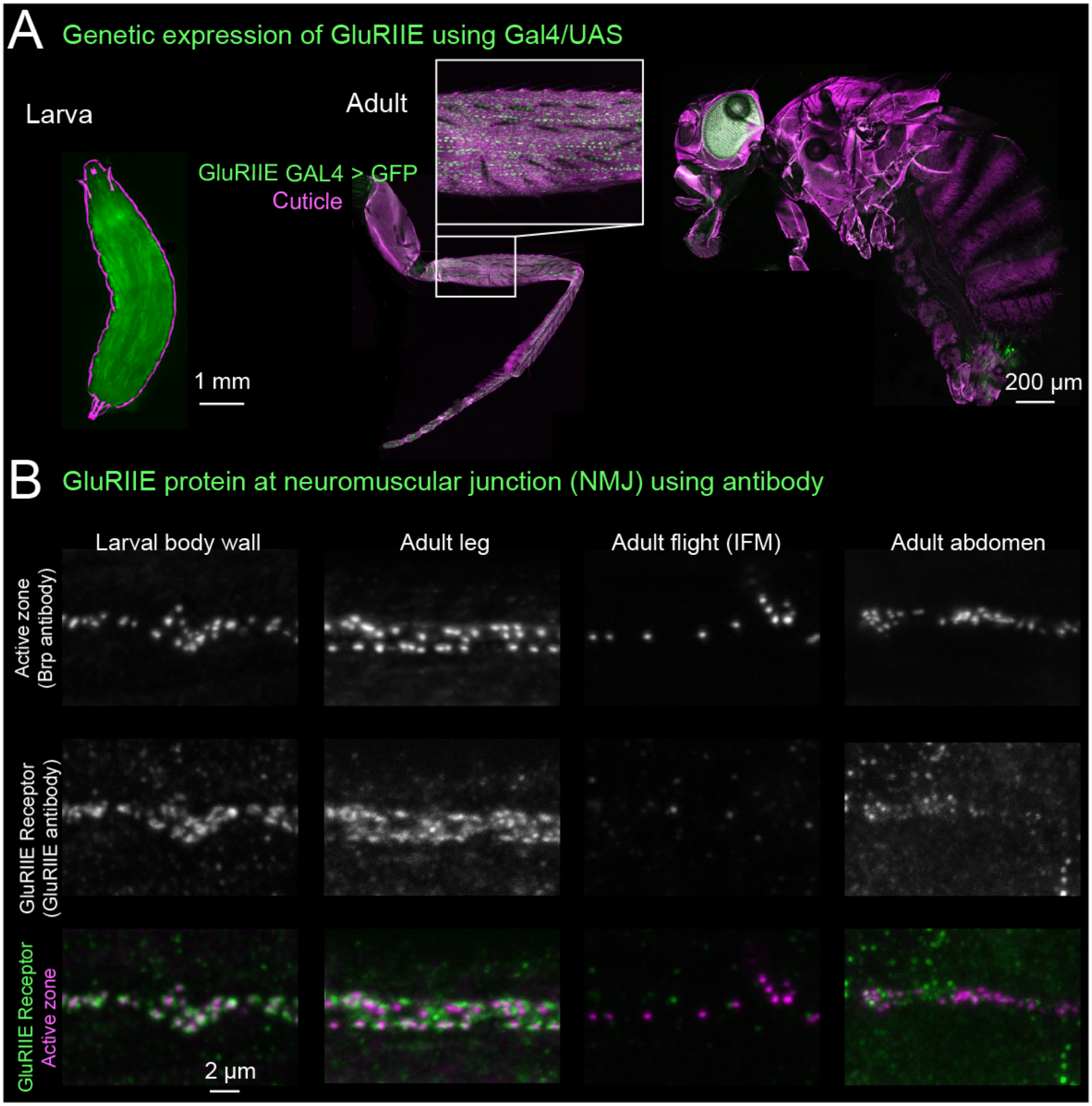


**Figure s8.** (A) Representative confocal images showing GFP expression driven by a GAL4 reporter line (GluRIIE-GAL4, green) in larval and adult muscles. (B) Representative images showing antibody staining of GluRIIE protein (green) at larval and adult neuromuscular junctions (magenta).

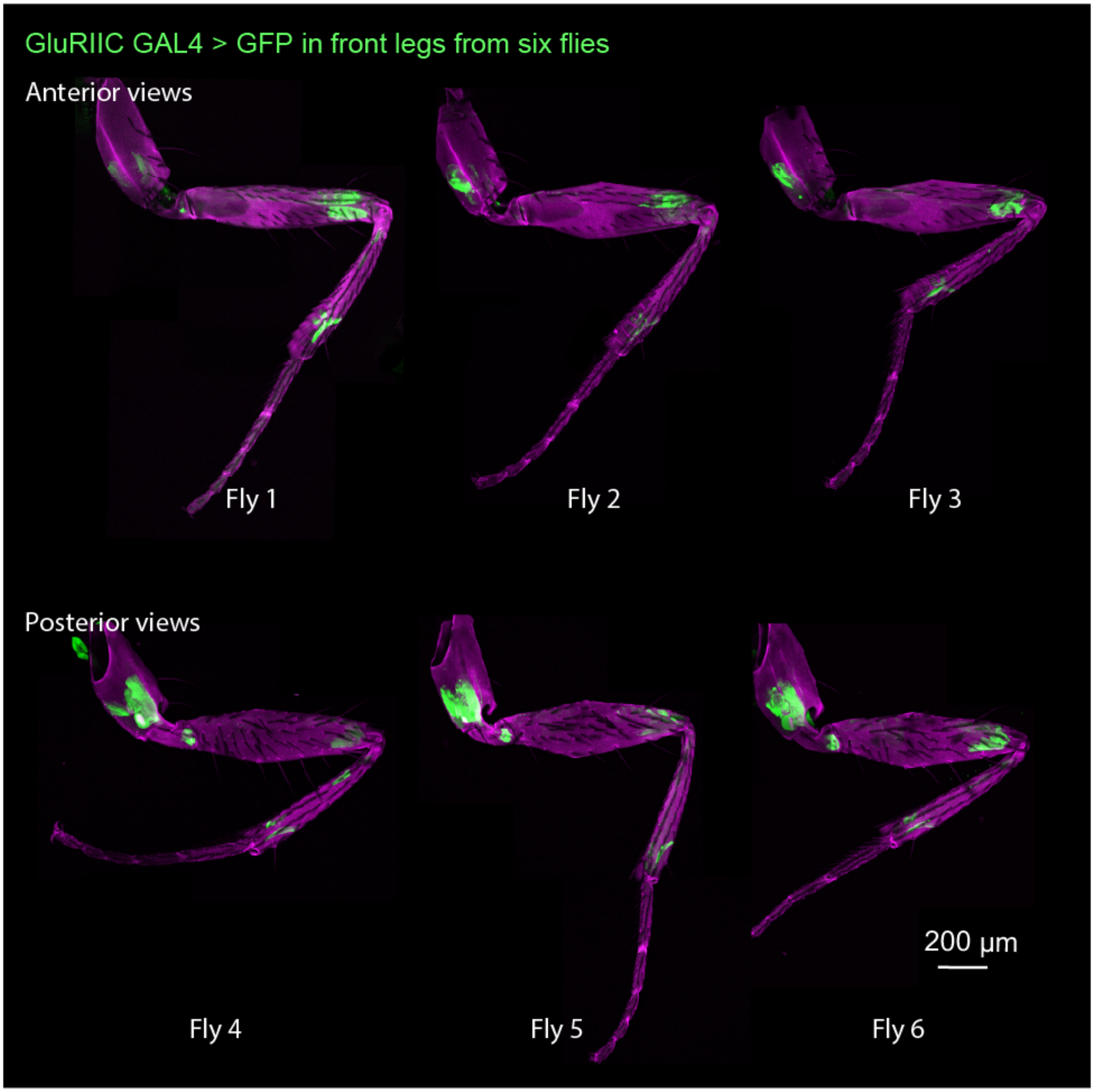


**Figure s9.** Representative confocal images showing GFP expression driven by a GAL4 reporter line (GluRIIC-GAL4, green) in six front legs from six flies. Anterior views (top), posterior views (bottom).

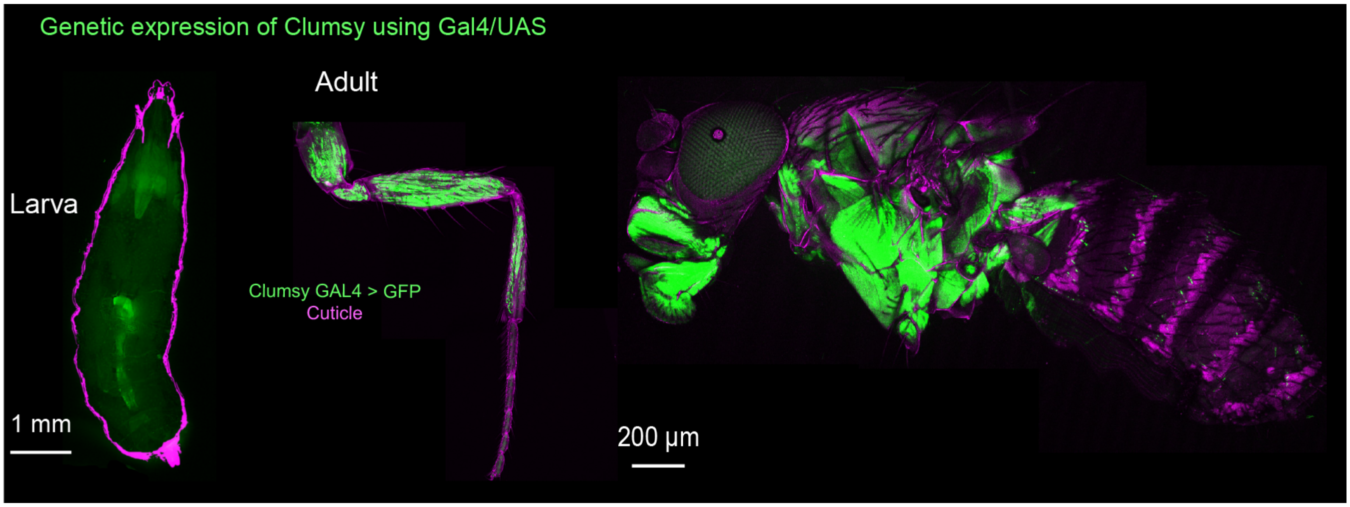


**Figure s10.** Representative confocal images showing GFP expression driven by a GAL4 reporter line (Clumsy-GAL4, green) in adult but not larval muscles. Note that the Clumsy trojan GAL4 has a 3xP3-GFP marker left from its construction that gives faint glial GFP expression in the brain, vnc, and gut that can be seen in the larva. Larval expression was not detected when we screened the line with a UAS-mCherry reporter. Adult muscle expression is consistent with both fluorescent reporters.

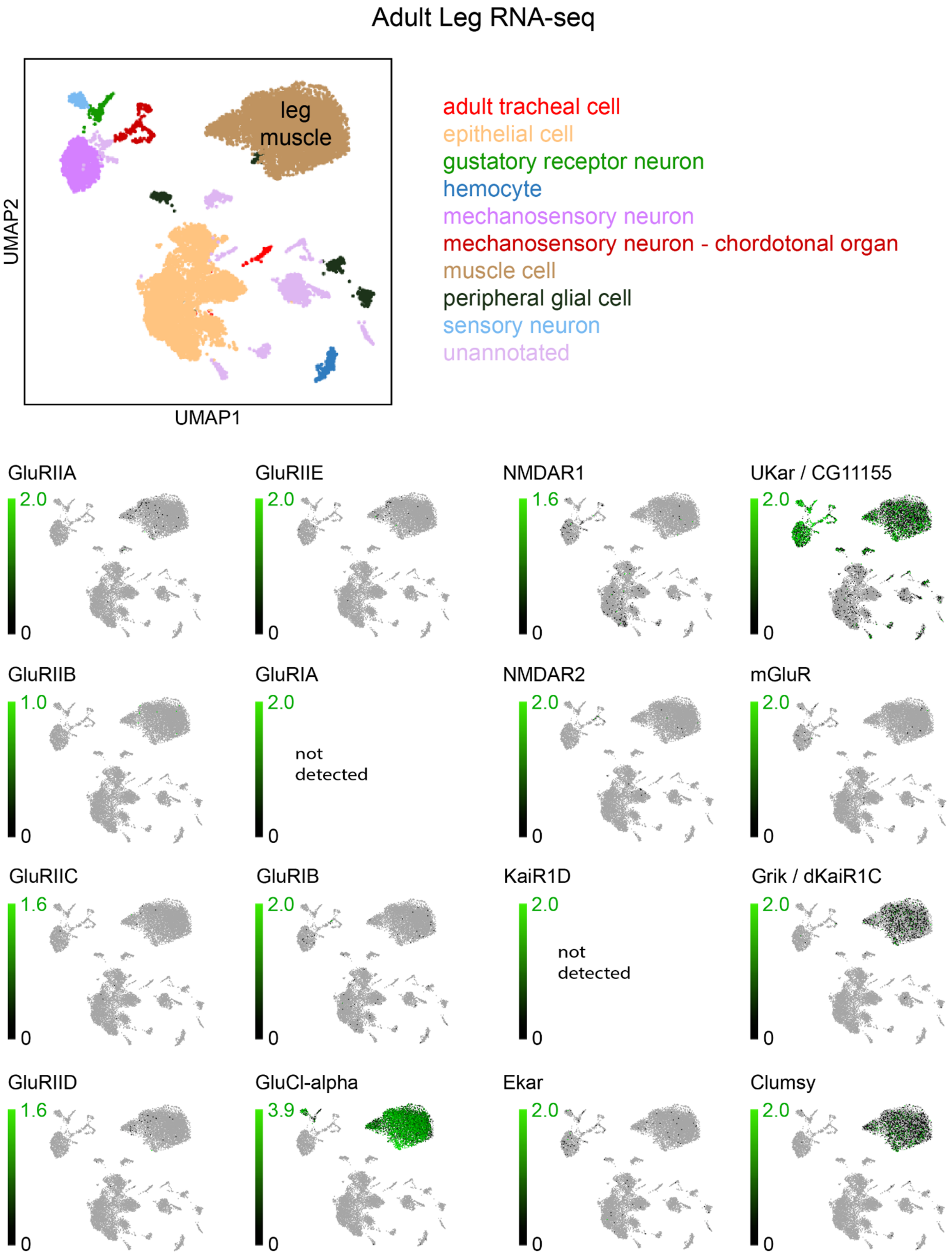


**Figure s11.** RNA-Seq analysis of glutamate receptors in all tissues of the adult leg (Li et al., 2022).

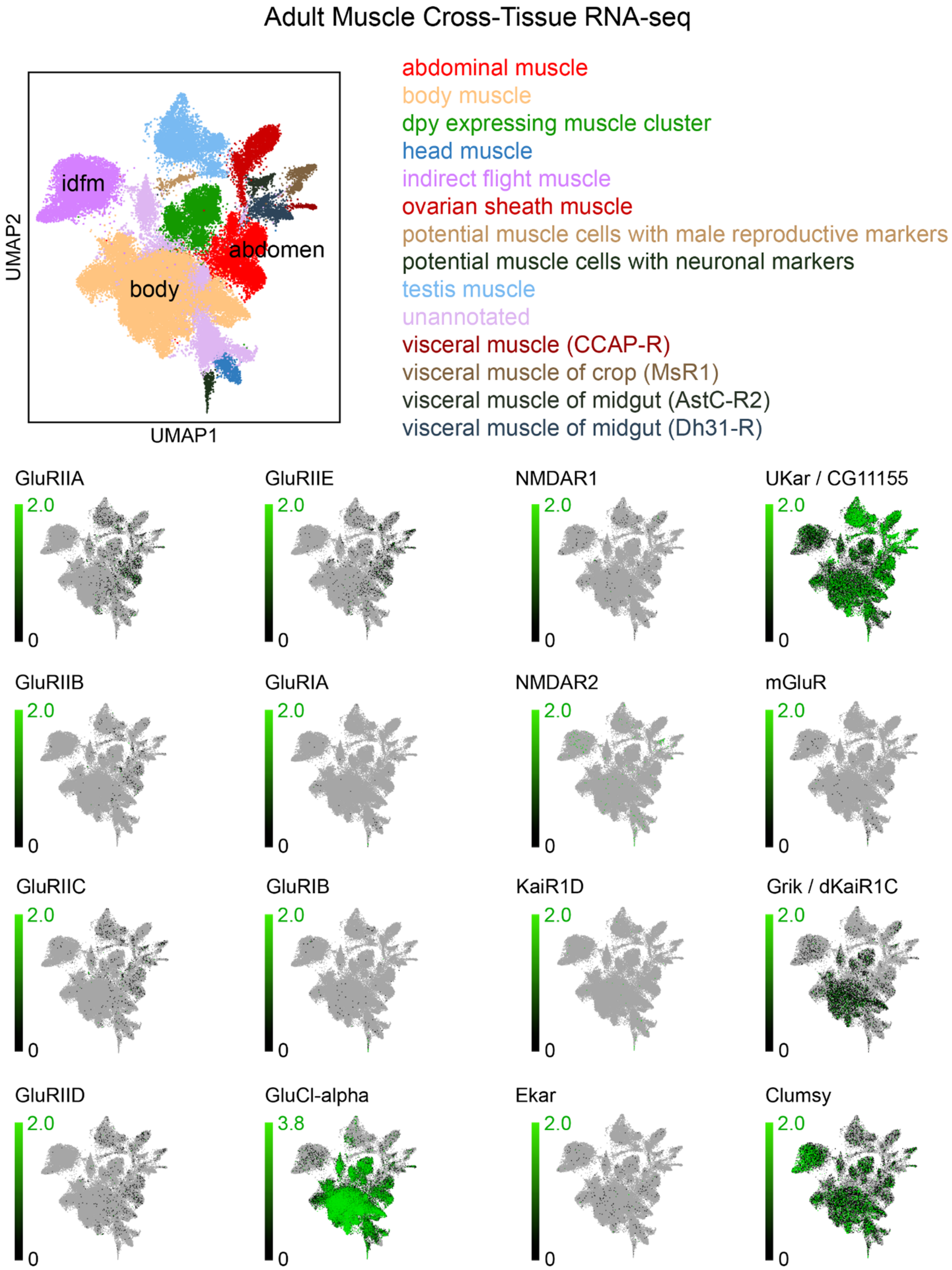


**Figure s12.** RNA-Seq analysis of glutamate receptors in all adult muscles (Li et al., 2022).

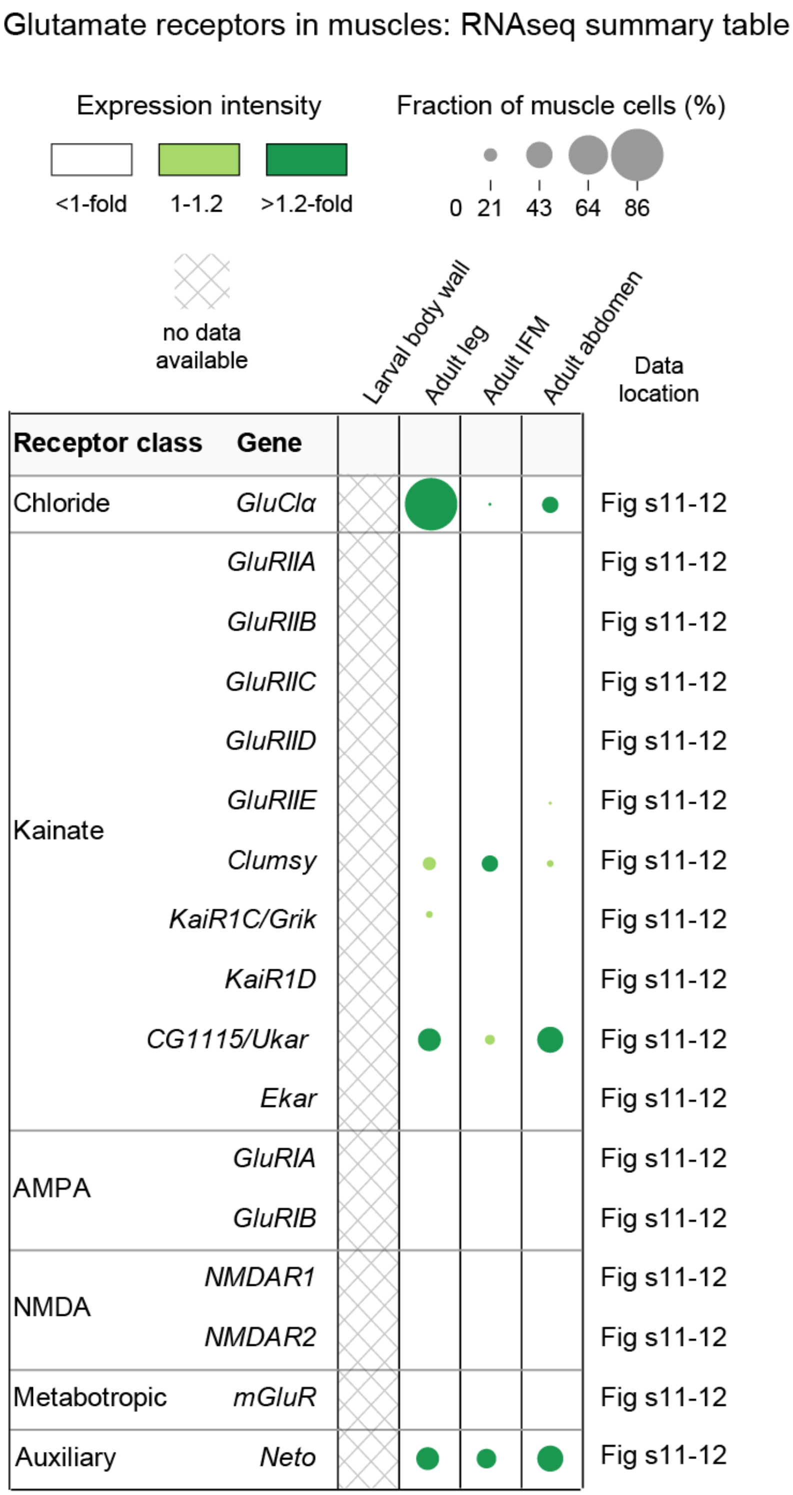


**Figure s13.** Table summarizing RNA-Seq data of glutamate receptors in adult muscles (data from Li et al., 2022).

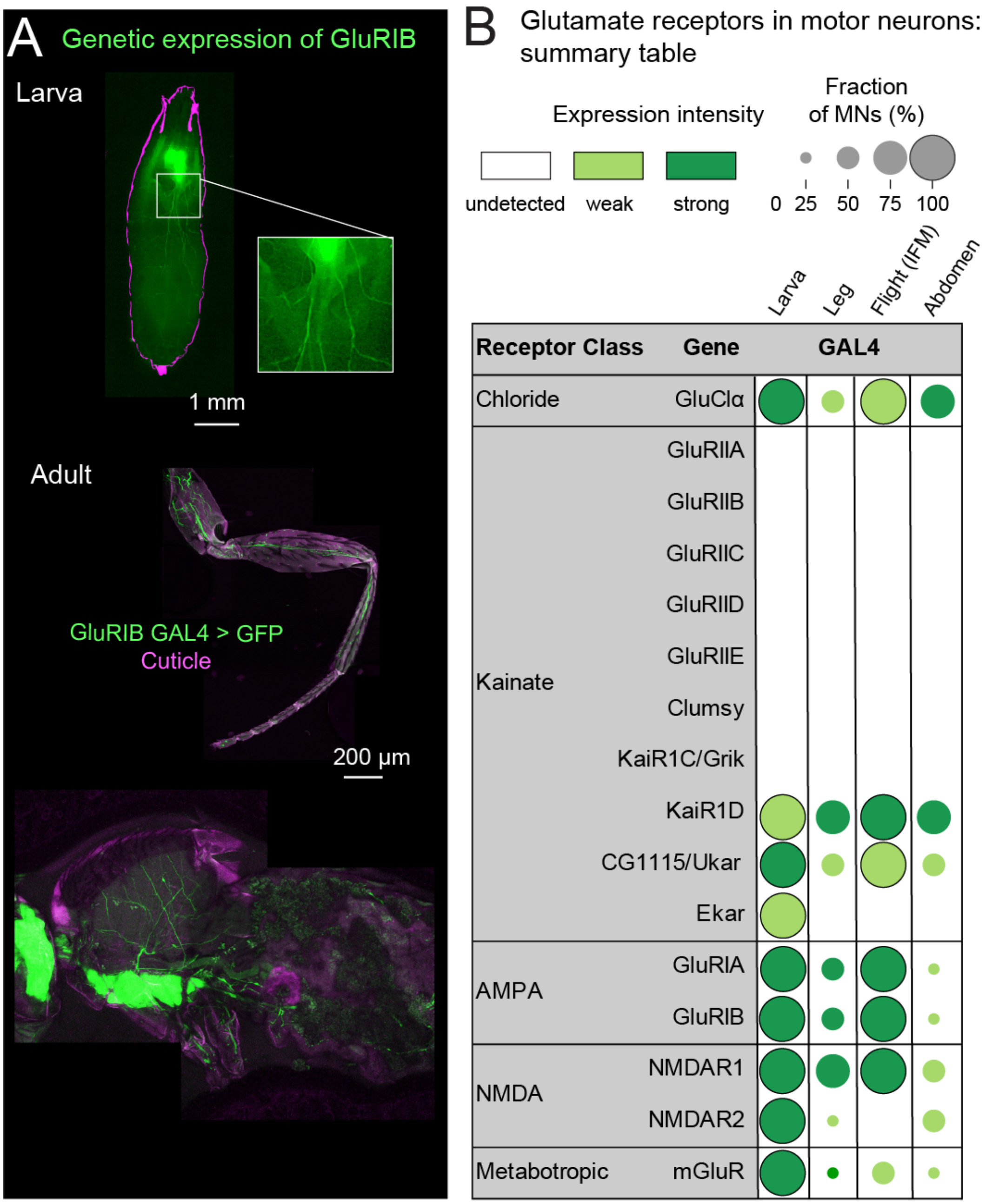


**Figure s14.** (A) Representative images showing expression of one example glutamate receptor subunit (GluRIB-GAL4, green) in larval and adult fly neurons. Motor neurons are visible as axons leaving the central nervous system. (B) Summary table of Gal4 expression in motor neurons.

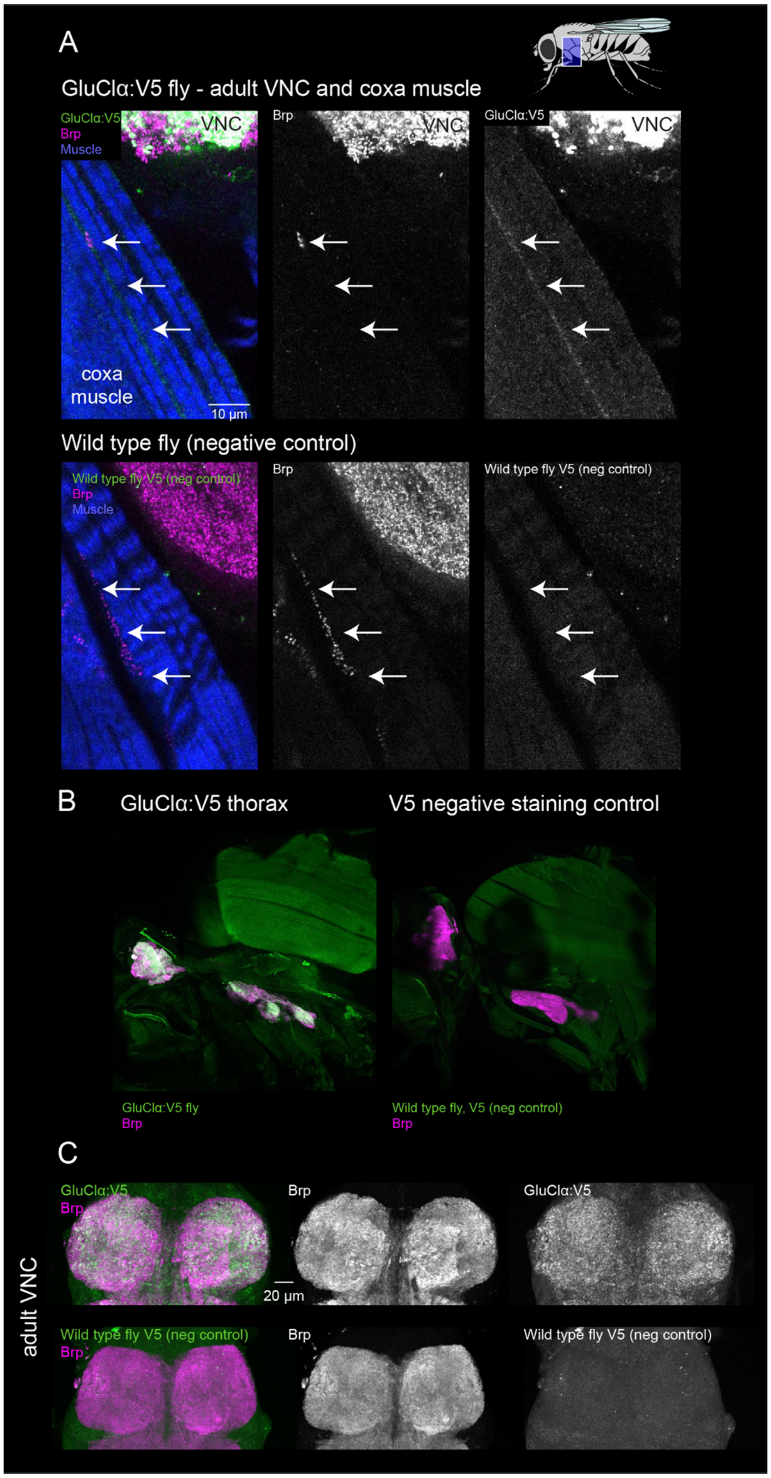


**Figure s15.** Validation of GluClα:V5 staining, showing positive controls (V5 staining in GluClα:V5 tagged flies) and negative controls (V5 staining in wild type flies, no tag). (A) Ventral nerve cord with nearby leg coxa muscle. Anti-V5 (green) localizes to the extrasynaptic muscle space (arrows) in the top fly (GluClα:V5) but not the bottom fly (wild type) (B) V5 staining positive and negative controls in the fly thorax. (C) V5 staining positive and negative controls in central nervous system. Antibody against V5 (green), antibody against Brp (magenta), and muscle (phalloidin, blue).
